# Integrated Analyses of Multi-omic Data Derived from Paired Primary Lung Cancer and Brain Metastasis Reveals the Metabolic Vulnerability as a Novel Therapeutic Target

**DOI:** 10.1101/2024.01.02.573855

**Authors:** Hao Duan, Jianlan Ren, Shiyou Wei, Chuan Li, Zhenning Wang, Meichen Li, Zhi Wei, Zhenyu Yang, Yu Liu, Yuan Xie, Suwen Wu, Wanming Hu, Chengcheng Guo, Xiangheng Zhang, Lun Liang, Chengwei Yu, Yanhao Mou, Yu Jiang, Houde Li, Eric Sugarman, Rebecca A. Deek, Zexin Chen, Likun Chen, Yaohui Chen, Maojin Yao, Lunxu Liu, Gao Zhang, Yonggao Mou

**Author notes:** These authors contributed equally (Hao Duan, Jianlan Ren, Shiyou Wei, Chuan Li, Zhenning Wang, Meichen Li). Correspondence to (Z.W.), (L.C.), (Y.C.), (M.Y.), (L.L.), (G.Z.) and (Y.M.).

## Abstract

Lung cancer brain metastases (LC-BrMs) are frequently associated with dismal mortality rates in patients with lung cancer; however, standard of care therapies for LC-BrMs are still limited in their efficacy. A deep understanding of molecular mechanisms and tumor microenvironment of LC-BrMs will provide us with new insights into developing novel therapeutics for treating patients with LC-BrMs. Here, we performed integrated analyses of genomic, transcriptomic, proteomic and metabolomic data which were derived from a total number of 174 patients with paired and unpaired primary lung cancer and LC-BrM, spanning four published and two newly generated patient cohorts on both bulk and single cell levels. We uncovered that LC-BrMs exhibited significantly higher intra-tumor heterogeneity. We also observed that mutations in a subset of genes were almost always shared by both primary lung cancers and LC-BrM lesions, including *TTN, TP53, MUC16, LRP1B, RYR2, and EGFR*. In addition, the genome-wide landscape of somatic copy number alterations was similar between primary lung cancers and LC-BrM lesions. Nevertheless, several regions of focal amplification were significantly enriched in LC-BrMs, including 5p15.33 and 20q13.33. Intriguingly, integrated analyses of transcriptomic, proteomic and metabolomic data revealed mitochondrial-specific metabolism was activated but tumor immune microenvironment was suppressed in LC-BrMs. Subsequently, we validated our results by conducting real-time quantitative reverse transcription PCR experiments, immunohistochemistry and multiplexed immunofluorescence staining of patients’ paired tumor specimens. Patients with a higher expression of mitochondrial metabolism genes but a lower expression of immune genes in their LC-BrM lesions tended to have a worse survival outcome. Therapeutically, targeting oxidative phosphorylation with gamitrinib in patient-derived organoids specific to LC-BrMs induced apoptosis and inhibited cell proliferation. The combination of gamitrinib plus anti-PD-1 immunotherapy significantly improved survival of mice bearing LC-BrMs. In conclusion, our findings not only provide comprehensive and integrated perspectives of molecular underpinnings of LC-BrMs but also contribute to the development of a potential, rationale-based combinatorial therapeutic strategy with the goal of translating it into clinical trials for patients with LC-BrMs.

## Introduction

Lung cancer is the second most commonly diagnosed cancer and one of primary causes of cancer-related mortality, with an estimated 2.2 million of new lung cancer cases and 1.8 million of patients’ deaths worldwide in 2020. This represents 11.4% of cancers diagnosed and 18.0% of cancer-related deaths(Sung et al., 2021). Lung cancer brain metastases (LC-BrMs) is one of the most frequent complications in patients with lung cancer(Riihimaki et al., 2014). As much as 20% of patients with lung cancer present with LC-BrMs at diagnosis and 50% at relapse(Siegel et al., 2018). Although cancer therapies have improved and patients tend to live longer with their primary tumors, the incidence of LC-BrMs has been rapidly increasing(Siegel et al., 2017).

LC-BrMs frequently causes mortality but standard of care therapies for LC-BrMs are still limited in their efficacy. Current treatment approaches toward LC-BrMs often utilize local therapy, such as radiation and/or resection in combination with systemic therapy, comprised of chemotherapy, targeted therapy and cancer immunotherapy. The blood-brain-barrier (BBB) selectively separates the brain from general circulation and this mechanism remains a significant barrier to effectively targeting BrMs with systemic therapy(Lowery and Yu, 2017). Present standard care of lung adenocarcinoma involves molecular testing of *epidermal growth factor receptor* (*EGFR*) and *anaplastic lymphoma kinase* (*ALK*) at the initial diagnosis due to multiple targeted therapies which have already been approved for this disease. The incidence of LC-BrMs among patients whose tumors harbor a mutation in *EGFR* or *ALK* is approximately half. Although brain is permeable for second or third-generation tyrosine kinase inhibitors (TKIs) targeting mutations in *EGFR* or *ALK*, a number of patients still have poor intracranial responses. Most patients exhibited either intrinsic or acquired resistance over time(Hida et al., 2017; Shaw et al., 2017; Solomon et al., 2018; Yang et al., 2015; Yang et al., 2016). This indicates additional genomic and non-genomic events may play a major role in promoting and sustaining BrMs.

In a study conducted by Brastianos *et al*., the analysis of whole-exome sequencing (WES) data derived from 86 “trios” of patient-matched pan-cancer BrMs, in comparison with primary tumors of various cancer types and normal blood samples demonstrated a pattern of branched evolution. They found that primary tumors and BrMs shared a common ancestor, yet BrMs possessed additional oncogenic alterations, which were not detected in up to 53% of primary tumors(Brastianos et al., 2015). Recently, Shih *et al*. characterized the genomic landscape of BrMs from 73 patients with lung adenocarcinoma via WES, and discovered increases in amplification frequencies of *MYC*, *YAP1* and *MMP13*(Shih et al., 2020).

Unlike many organs in which extracranial metastases develop, the brain does not share a relatively similar composition at the cellular level with the organ in which the primary tumor originated(Zhang et al., 2013). Instead, the brain itself consists of a wide range of cellular types, including neurons, astrocytes, microglia and etc., which are uniformly absent in the extracranial organs from which primary cancers originated. The multifaceted cellular process by which cancer cells adapt to this highly specialized tumor microenvironment may also involve additional steps of selecting and enriching modifications that occur genetically and epigenetically. By performing multi-omic analyses of a cohort of the lung, breast, and renal cell carcinomas consisting of BrMs and matched primary or extracranial metastatic tissues, Fukumura *et al*. demonstrated that oxidative phosphorylation (OXPHOS) was prominently upregulated in BrMs and inhibition of OXPHOS alone impaired BrMs of breast cancer cell lines in preclinical models(Fukumura et al., 2021). Indeed, metabolic reprogramming is a hallmark of cancer(Hanahan, 2022). Although tumor cells prefer to generate energy via glycolysis even in the presence of oxygen, they would reprogram the metabolic network towards OXPHOS, which provides tumor cells with a new source of energy to facilitate their survival under stress stimuli and adaptation to a hostile environment. In the past decade, several inhibitors have been developed to target OXPHOS in tumor cells, including phenformin(Birsoy et al., 2014; Masoud et al., 2020), IACS-010759(Molina et al., 2018), gamitrinib(Chae et al., 2012; Chae et al., 2013; Kang et al., 2009; Wei et al., 2022; Zhang et al., 2016), and several others, among which gamitrinib has demonstrated its selectivity and specificity. It is worth noting that gamitrinib has entered phase I clinical trial as a monotherapy to treat patients with advanced tumors (ClinicalTrials.gov Identifier: NCT04827810).

Considering the selective pressure of brain-resident cells acting on metastatic cells, emerging evidence has suggested a new avenue for therapeutic intervention of BrMs that is to target the crosstalk between cancer cells and the microenvironment(Fischer et al., 2019). Single-cell RNA sequencing (scRNAseq) of different metastatic lesions including BrMs derived from lung adenocarcinoma demonstrated vibrant cell-population dynamics and molecular interactions between the tumor, stromal, and immune compartments, which creates a pro-tumoral and immunosuppressive microenvironment(Kim et al., 2020). The aforementioned study performed by Fukumura *et al*. also documented a decrease in cellular immunity in BrMs relative to patient-matched primary or extracranial counterparts(Fukumura et al., 2021). Zhang *et al*. also found that Immunosuppressive tumor microenvironment in brain metastasis (Zhang et al. 2022). Recent studies showed that intracranial and extracranial benefits to immune checkpoint blockade monotherapy or in combination use with chemotherapy were near identical in patients with lung cancer(Gandhi et al., 2018; Goldberg et al., 2020; Rittmeyer et al., 2017). The hazard ratio (HR) for overall survival (OS) with atezolizumab vs docetaxel in the OAK study was 0.73, 0.74 and 0.54 for the overall population, patients without BrMs, and patients with BrMs, respectively(Rittmeyer et al., 2017). Similarly, the HR for OS in the KEYNOTE-189 study of pembrolizumab and chemotherapy vs chemotherapy alone was 0.49 for the overall population, 0.36 for patients without BrMs, and 0.53 for patients with BrMs (Gandhi et al., 2018). Understanding the tumor microenvironment of LC-BrMs will further improve the efficiency of current immunotherapies.

In order to comprehensively reveal the genetic alterations, transcriptomic dynamics, proteomic modifications and specialized microenvironment in LC-BrMs, we undertook a holistic approach by conducting integrated analyses of primary lung–brain metastasis pairs at genomic, transcriptional, proteomic and metabolomic levels using WES, bulk RNAseq, proteomics, reverse phase protein array (RPPA), and metabolomics. The multi-omic results were further strengthened by scRNAseq, mitochondrial DNA (mtDNA) content analysis, multiplex immunofluorescence (mIF) and immunohistochemistry (IHC) staining. Moreover, patient-derived organoids (PDOs) of LC-BrMs and a mouse LC-BrM model were used to investigate the therapeutic efficacy of gamitrinib and the combination of gamitrinib plus anti-PD-1 antibody on LC-BrMs, respectively. This study has not only uncovered new molecular characteristics of LC-BrMs but also nominated a combination approach by targeting OXPHOS and reinvigorating the tumor immune microenvironment.

## Results

### Patient cohorts

In this study, we assembled the largest cohort of patients with paired primary lung cancer and BrM lesions, for all of which multi-omic data were available. We computationally analyzed WES, bulk RNAseq and RPPA data available from three published datasets(Brastianos et al., 2015; Fukumura et al., 2021; Shih et al., 2020) (**Fig. 1a**). In addition, we performed WES, bulk RNAseq, proteomics and metabolomics of a newly generated cohort of patients presented at Sun Yat-sen University Cancer Center (SYSUCC) (**Fig. 1a**), among which RNAseq data of 24 patients was published in our previous study(Li et al., 2022). Overall, a total of 119 patients with paired primary lung cancer and BrM samples were included for genomic analysis; 56 patients were included for transcriptomic analysis; 16 patients were included for proteomic analysis (4 for proteomics and 12 for RPPA); and 4 patients were included for metabolomic analysis.

**Figure 1.**
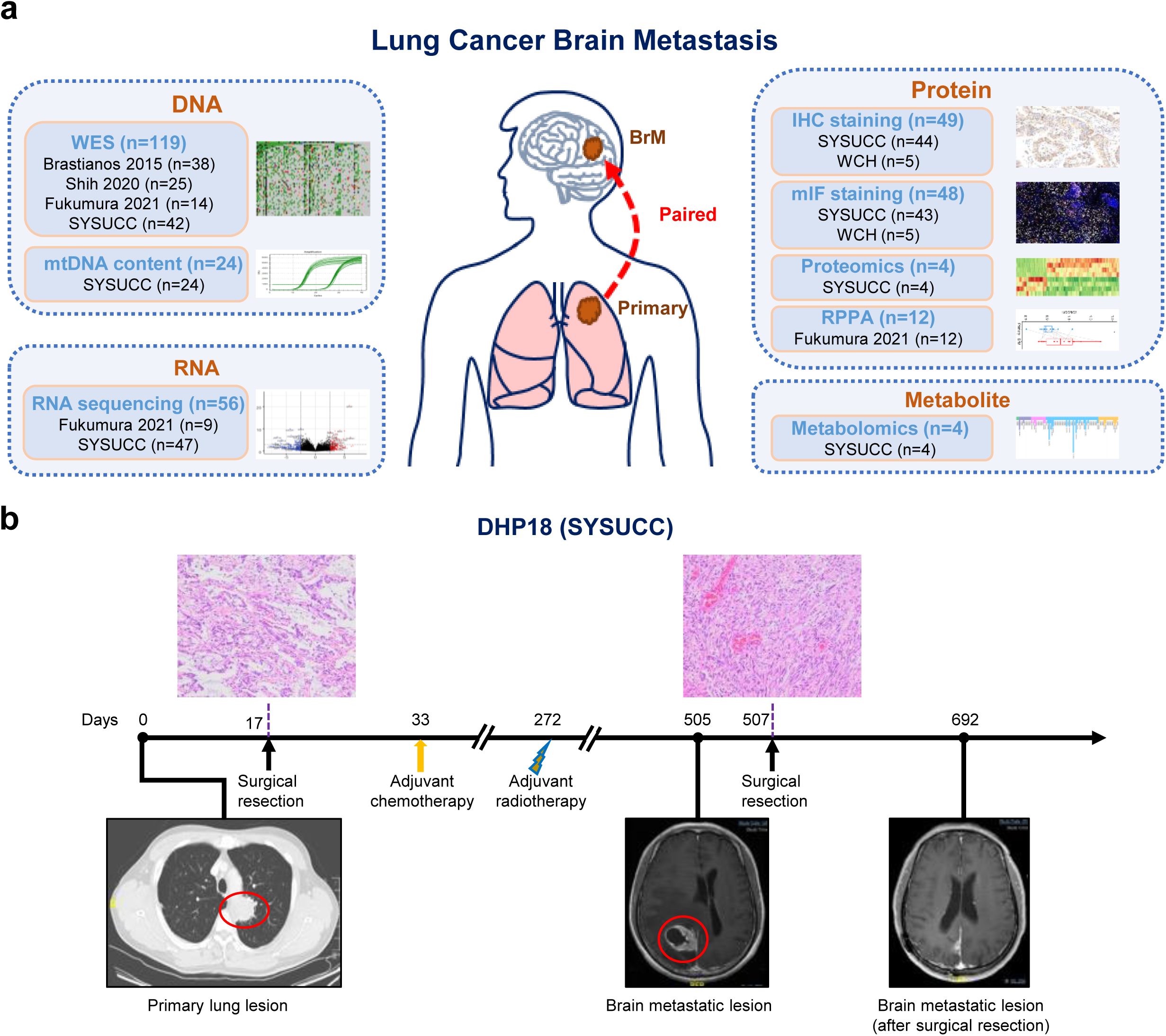
Study workflow, overview of patients and samples, and cohort characteristics. **a**. Overview of patient cohorts and various experimental platforms. **b**. A representative patient with primary lung cancer (DHP18) who later developed brain metastasis. A primary lung lesion was detected by CT scan and surgical resection was performed to obtain the primary lung cancer sample. However, one year after the surgery, a new brain metastasis lesion was identified through MRI scan. The surgical resection was then performed to remove the brain metastatic lesion.

To validate the results of multi-omic data, an independent cohort of scRNAseq(Kim et al., 2020) data that consisted of 45,149 single cells derived from 15 primary lung cancers and of 29,060 single cells derived from 10 BrM lesions was investigated. Additionally, we further assembled two patient cohorts with paired primary lung cancer and BrM lesions from SYSUCC and West China Hospital (WCH) which were used for IHC (n=49) and mIF (n=48) staining (**Fig. 1a**). Besides, 24 patients from the SYSUCC cohort were included for mtDNA content analysis. Clinicopathological characteristics of all cohorts and analytical approaches of each sample were summarized in **Table 1, Table S1** and **Table S2**, respectively. The analytic pipelines of WES, RNAseq, scRNAseq, proteomics and metabolomics data were listed along with the tools that were used in each step in **Supplemental Fig. 1**.

**Table 1.**
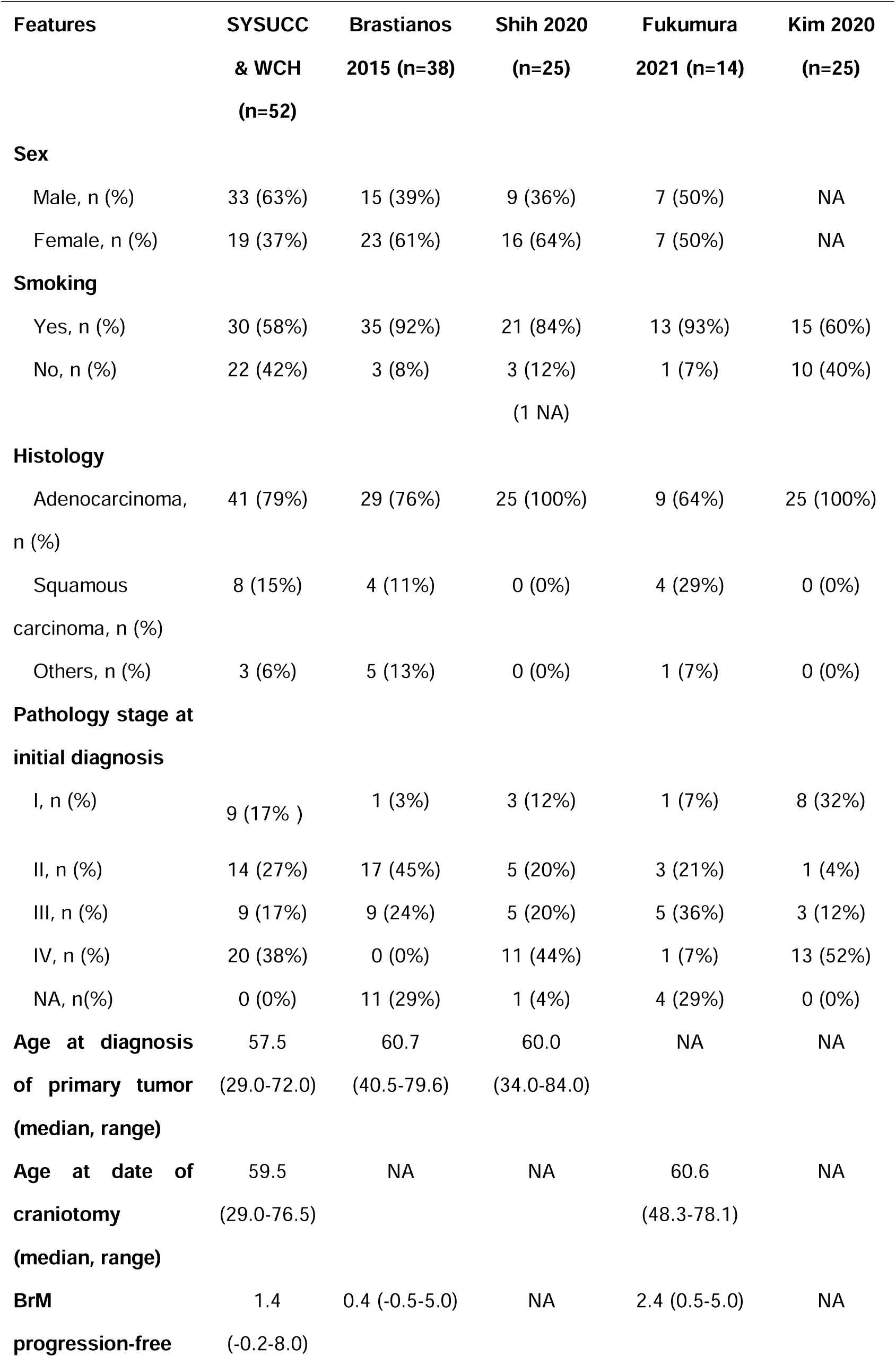
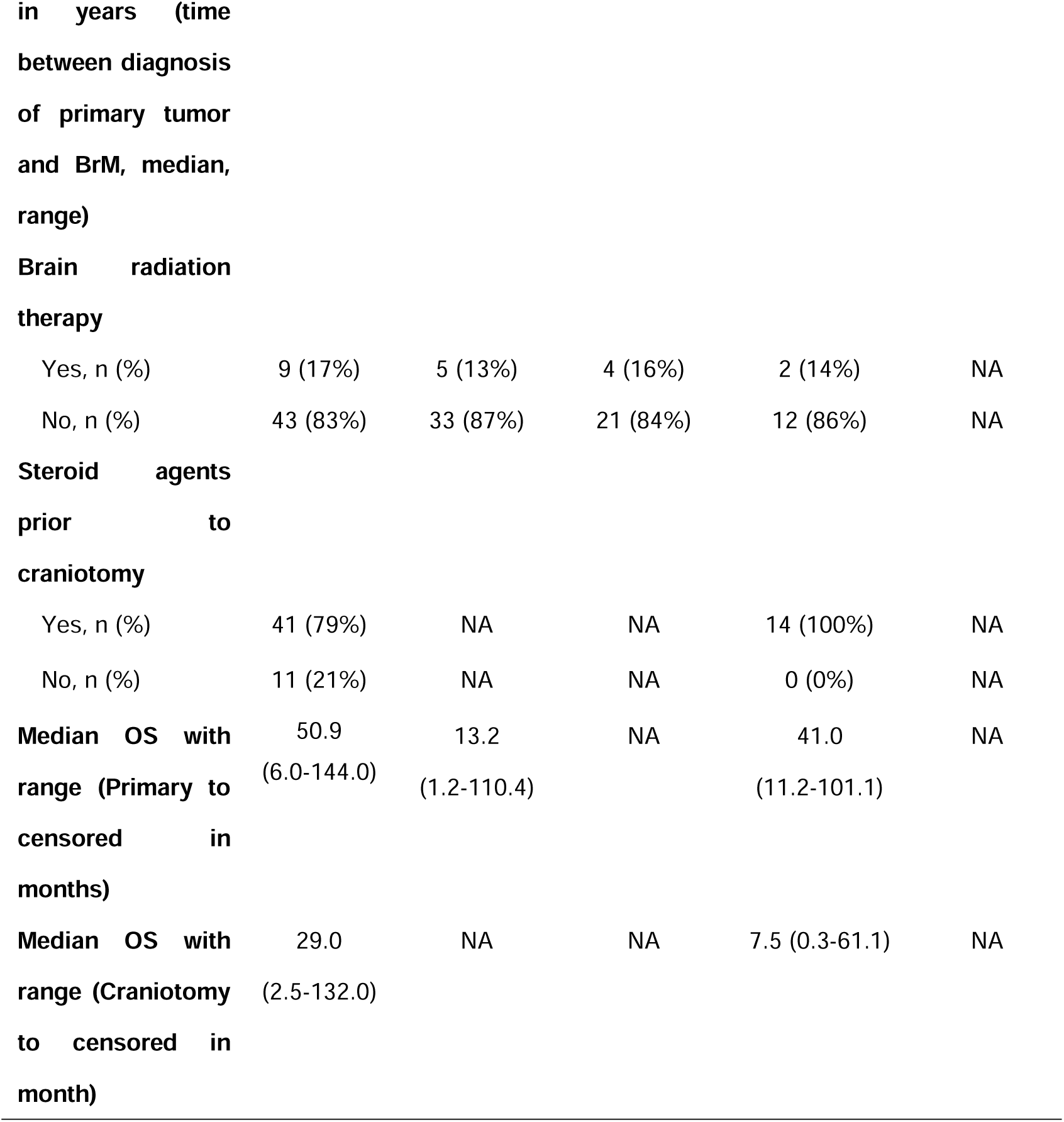
Clinicopathological characteristics of included cohorts.

Took the 51-year-old male, patient DHP18 from the SYSUCC cohort, as an example: a lung mass at the upper left lobe was identified by computerized tomography (CT) scan on day 0 and confirmed as pathologic stage III (T4N0M0) lung poorly differentiated adenocarcinoma on day 17. Although adjuvant chemotherapy was administrated after lobectomy on day 33 and radiotherapy was administrated for local recurrence on day 272, BrM was detected on day 505 and subsequently resected on day 507 (**Fig. 1b**). H&E staining was performed to confirm the pathological diagnosis of primary lung cancer and BrM lesion.

To our best knowledge, our endeavor represented the largest integrated analyses of multi-omic data that comprehensively depicted BrM lesions derived from primary lung cancer.

### Frequencies of somatic alterations in primary lung cancers and LC-BrMs

We evaluated how stability, acquisition and loss of somatic mutations would affect the metastasis of primary lung cancers to the brain. We used the case/control somatic alteration analysis to compare LC-BrMs to primary lung cancers and presented the top 25 ranked mutations for paired primary lung cancers and LC-BrMs, respectively (**Fig. 2a, b**). We observed that mutations in a subset of genes were almost always shared by both primary lung cancers and BrM lesions, including *TP53* (42%, 49%), *MUC16* (46%, 45%), *LRP1B* (43%, 42%), *TTN*(62%, 64%), *OBSCN* (26%, 25%), *FAT3* (25%, 26%) and *EGFR*(22%, 20%), which were commonly mutated in lung cancers(Cancer Genome Atlas Research, 2014) (**Fig. 3a, b**, **Supplemental Fig. 3a**). Our analysis also revealed mutations in *TTN, TP53, MUC16, LRP1B, RYR2, EGFR and BRAF* were shared between primary lung cancers and LC-BrMs. These findings were consistent with previous reports, showing that these somatic mutations were frequently identified in lung adenocarcinoma(Cancer Genome Atlas Research, 2014). The forest plot of overall frequency of somatic alterations in primary lung cancers and LC-BrMs was shown in **Supplemental Fig.3b**. The higher frequency (i.e., above the median frequency cutoff values) of each of the ten genes – *UTRN, DGKH, IL17RA, ITGA6, CUL1, DDX50, MTFR1, EXOC7, NUP188* and *KDM5B* was significantly associated with primary lung cancer, while odds ratios (OR) of *MUC2* and *ELAVL2* were significantly less than 1 in the BrM cohort. Our data suggested that *MUC2* and *ELAVL2* might be the driver genes which were specifically mutated in LC-BrMs (**Supplemental Fig. 3b**).

**Figure 2.**
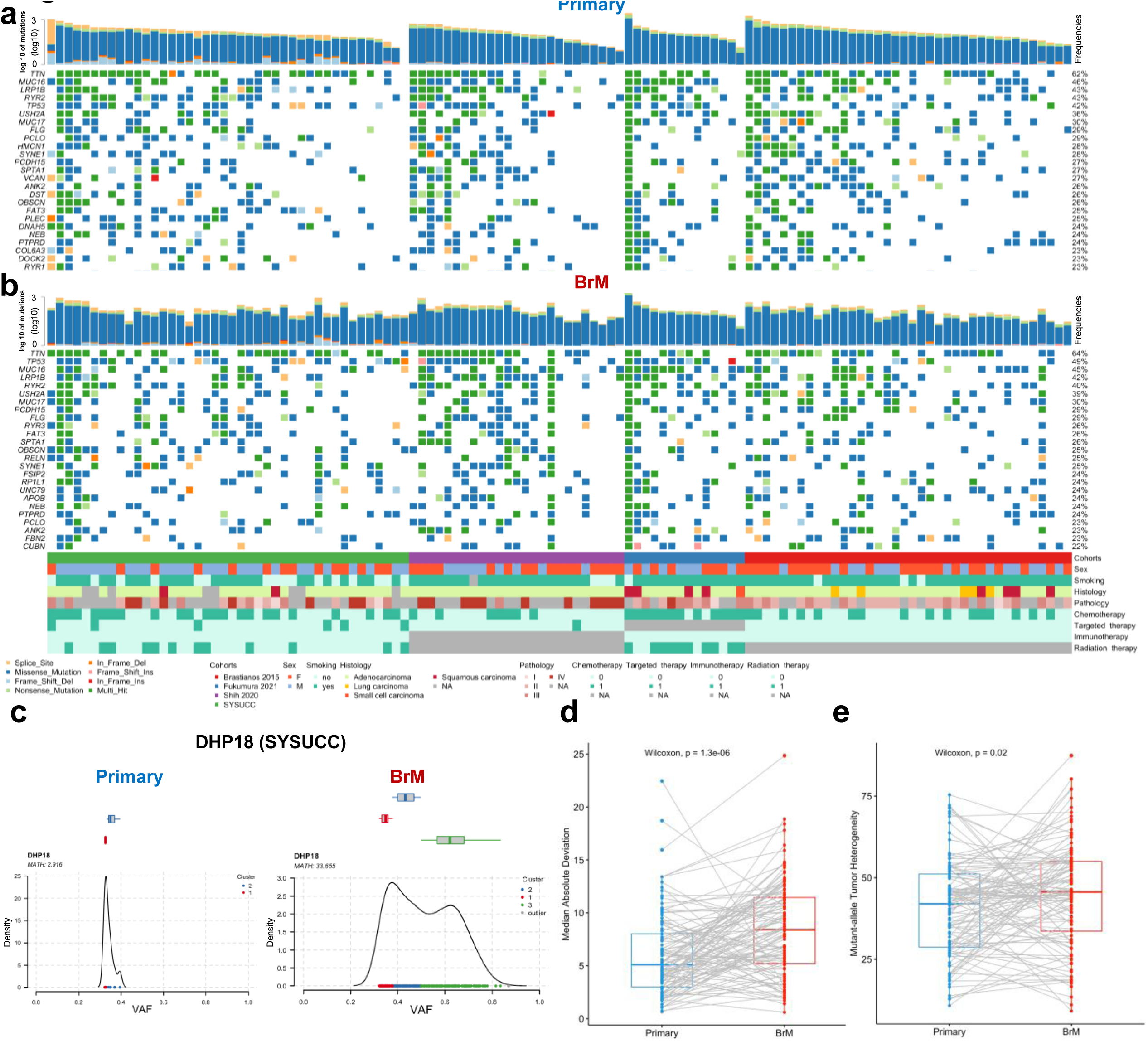
Oncoplots showing the 25 most frequently mutated genes in paired primary lung cancer and brain metastasis (BrM) specimens. **a and b.** Recurrently mutated oncogenic driver genes identified by Mutect2 in primary lung cancers (**a**) and BrMs (**b**). Every column represented a single patient stratified by cohort and ordered from left to right by the number of oncogenic driver genes identified in BrM samples. The variant type was depicted by its color. **c**. Cluster plots of the primary lung cancer (left) and BrM sample (right) derived from the representatrive patient DHP18. X-axis represented variant allele frequency, the top bar showed the number of clusters on top of each plot, and the math number was noted in the upper left corner. d and e. The box plot of median absolute deviation and mutant-allele tumor heterogeneity in paired primary lung cancers and brain metastasis lesions. The p-value was determined using pairwise two-sided Wilcoxon test.

**Figure 3.**
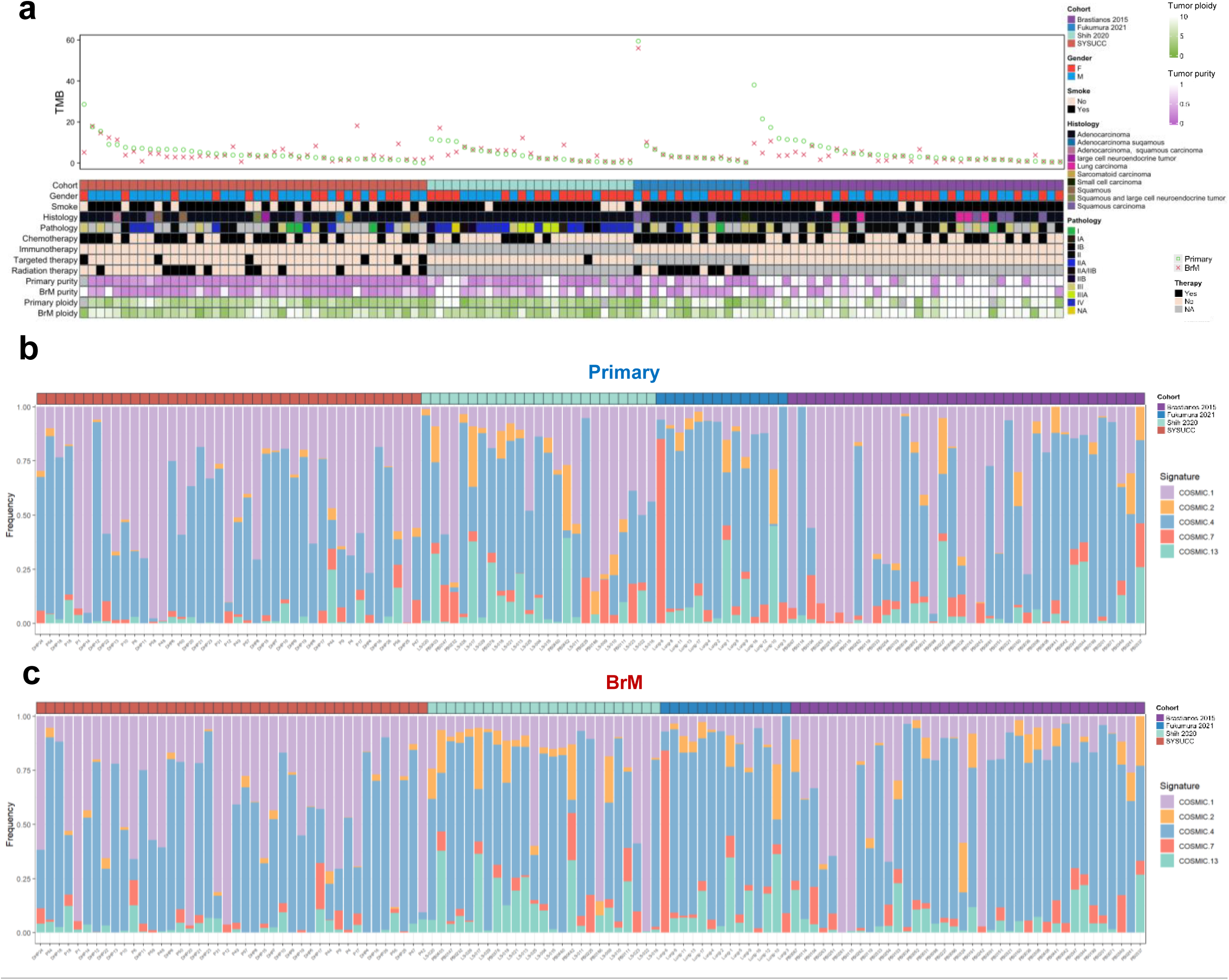
Tumor mutational burden and mutational signatures. **a.** Integrated analyses of four indepedent cohorts (n=119 patients) depicting TMB in each pair of primary lung cancer and brain metastasis (BrM) specimens according to cohorts, gender, smoking history, histological group, pathological level, different therapies, purity and ploidy. Each column represents a single patient with two tumor specimens scattered at two separate spaces. All tumor specimens were grouped by cohorts and ordered from left to right by decreasing mutation frequencies of BrMs. Green circle indicated primary lung cancer and the red fork indicated BrM. **b**. Percentage of mutational signature contribution in each tumor sample.

We also inferred intra-tumor heterogeneity (ITH) in primary lung cancers and BrM lesions by clustering variant allele frequencies (VAF)(Mroz and Rocco, 2013). The average of median absolute deviation of primary lung cancers was 4.92 as compared to 8.25 for BrM lesions, and mean mutant-allele tumor heterogeneity (MATH) was 35.15 for primary lung cancers compared to 42.32 for LC-BrMs. Differences between primary lung cancers and LC-BrMs were significant for both scores (p = 0.00019 and 0.036, respectively) (**Fig. 2d, e**), indicating that LC-BrMs exhibited a higher ITH compared to primary lung cancers. For example, the primary lung cancer derived from the patient DHP18 showed no separation of clones clustered at mean variant frequencies of ∼35% with a MATH score of 2.916, while the LC-BrM showed a clear separation of two clones clustered at ∼35% major clone and ∼65% minor clone with a MATH score of 33.655 (**Fig. 2c**). Two other pairs of primary lung cancers and LC-BrMs also showed similar results (**Supplemental Fig. 3e, f**). Together, the persistence of drivers and the paucity of consistent changes indicated that BrM lesions were characterized by a higher ITH compared to paired primary lung tumors.

### The somatic landscape of primary lung cancers and LC-BrMs

Subsequently, we analyzed WES data by calculating tumor mutational burden (TMB), including SNPs, indels, and CNVs, in order to understand general patterns of primary lung cancers and their paired BrM lesions. TMB exhibited by primary lung cancers and LC-BrMs were comparable to the previously reported findings from the TCGA-LUAD cohort containing 516 samples with median TMB of 7.78 (Ellrott et al., 2018) (**Supplemental Fig. 2a**). Median TMBs were 6.22 and 5.21 for primary lung cancers and BrM lesions, respectively, as defined by single-nucleotide variants and small insertions and deletions (indels) per megabase (Mb). The increase in TMB-SNPs exhibited by BrM occurred in 15 out of 119 patients, whereas the decrease in TMB occurred in 104 out of 119 patients (**Fig. 3a**). Next, we evaluated differences of TMB, SNP, INDEL, arm-level CNV and gene-level CNV between two cohorts (**Supplemental Fig. 2b**). There were no significant differences of TMB and SNP between primary and BrMs (Wilcoxon p-value = 0.2 and 0.34, respectively); however, INDELs TMB, arm-level CNVs TMB and gene-level CNVs TMB were significant different between primary lung cancers and LC-BrMs (Wilcoxon p-value = 0.063, 1.4e-08 and 1.4e-07, respectively).

These similarities and differences of somatic alterations in primary lung cancers and BrM also contributed to components of COSMIC mutational signatures. As expected, the signature activity was closely related to somatic mutations as we observed similar patterns of mutational signatures exhibited by primary lung cancers and LC-BrMs (**Fig. 3b, c**). Signature 1 - spontaneous or enzymatic deamination of 5-methylcytosine, signature 4 - exposure to tobacco, and signature 7 - UV exposure were nearly always the dominant signatures among all cohorts. Signature 2 - viral infection, retrotransposon jumping or to tissue inflammation and signature 13 - APOBEC C>G were dominantly detected in both Brastianos and Fukumara cohorts; however, the cosine similarities were all under 0.5 overall, which were not considered as significant. Interestingly, signature 2 is attributed to the activity of the APOBEC family of cytidine deaminases, which was found in 22 different cancer types but most commonly in cervical and bladder cancers.

### The landscape of somatic copy number alterations (SCNAs)

We next assessed SCNAs and found that the genome-wide landscape of SCNAs was similar between primary lung cancers and LC-BrMs (**Fig. 4a, b and Supplemental Fig. 4a-f**). Chromosome arm-level copy number events occurred with similar frequencies in all four cohorts. Across all four cohorts, our analysis revealed 20 arm-level gains and 31 arm-level losses exhibited by LC-BrMs as compared to 25 arm-level gains and 33 arm-level losses exhibited by primary lung cancers. Among these SCNA regions, 18 out of 20 gains and 28 out of 31 losses were shared by primary lung cancers and BrM lesions. Moreover, we identified peaks of 24 amplifications and 45 deletions in LC-BrMs compared to peaks of 29 amplifications and 57 deletions identified in primary lung cancers (FDR q-value < 0.25). Among these peaks, gains in 5p15.33, 7p11.2, 10q11.21, 11q13.1, 11q13.3, 12q15, 14q13.3 and 20q13.33 chromosomal arms were shared by primary lung cancers and BrM lesions, while losses in 1p36.21, 6p21.33, 6q25.3, 9p21.3 and 17p11.2 were shared by primary lung cancers and BrM lesions (**Supplemental Fig. 4a, b**). Among these shared peaks, the largest peak was an amplification of 20q13.33 with 149 genes affected. The narrowest peaks affecting a single locus included amplifications of 7p11.2, 11q13.1, 11q13.3, 12q15 and 14q13.3; and deletions of 6q25.3, 9p21.3 and 17p11.2. The three most common somatic copy number peaks occurred in 7p11.2 (26% vs. 37%), 12q15 (12% vs. 15%), and 11q13.3 (21% vs.12%) in both primary lung tumors and LC-BrMs (**Supplemental Fig. 4a, b**). We applied Genomic Identification of Significant Targets In Cancer (GISTIC), which is an established methodology(Mermel et al., 2011) to compute a SCNA positive selection score for each genomic location. This approach assessed amplitudes and frequencies of SCNAs across samples and identified regions with significantly recurrent SCNAs that likely resulted from a positive selection. The highest-ranked genes in both cohorts of primary lung cancer and LC-BrMs included *CCNL2*, *DVL1*, *ATAD3A*, *AGRN* and ISG15, where mutation patterns were clustered for the patients (**Supplemental Fig. 4e-f**).

**Figure 4.**
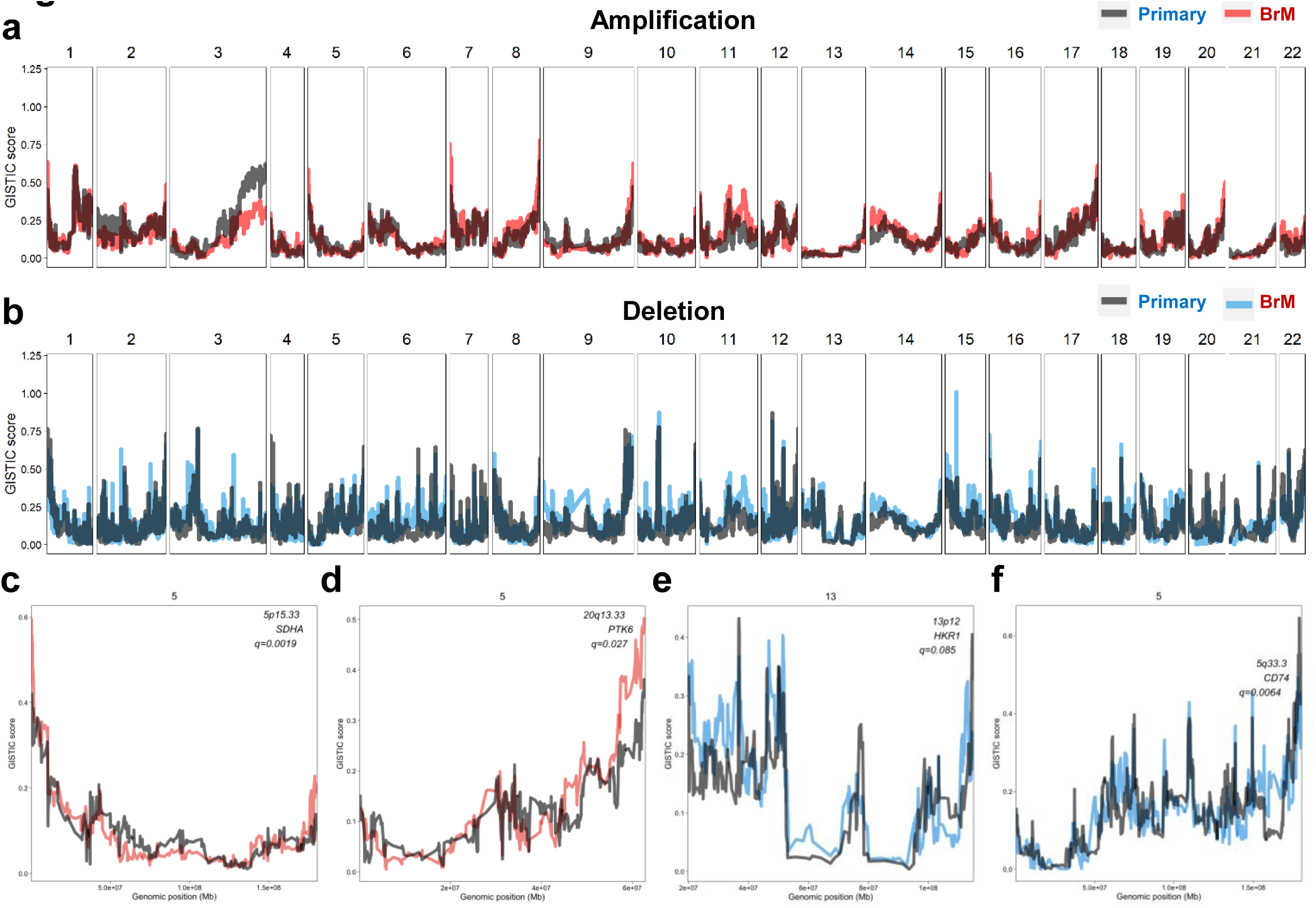
The landscape of somatic copy-number alteration in paired primary lung cancer and brain metastasis (BrM) specimens. **a and b.** GISTIC amplification (**a**) and deletion (**b**) plots of primary lung cancer (n=119) and BrM (n=119) samples. In the top figure, red line is the amplification in BrM, and grey line is amplification in primary; in the bot figure, blue line is the deletion in BrM, and grey line is deletion in primary. **c-f**. Four representative GISTIC plots showing regions of candidate driver genes of BrM compared to primary lung cancer.

Despite of broad similarities of copy-number landscapes between primary lung cancers and LC-BrMs, four distinct genomic regions with significantly different scores of positive selection were determined (**Fig. 4c-f**). We identified 52 out of 226 unique deletions and 5 out of 15 unique amplifications in BrM with q-value less than 0.1 (Supplemental materials). Two regions of unique focal amplifications that were significantly enriched in BrM (q-value = 0.0019 and 0.027, respectively), including: (1) 5p15.33 containing *SDHA, PDCD6, AHRR, CCDC127, PLEKHG4B,* and *LRRC14B*; (2) 20q13.33 containing *MYT1, PTK6* and etc. Two regions of unique focal deletions (q-value = 0.085 and 0.0064, respectively) include (1) 19q13.12 containing *HKR1* and (2) 5q33.3 containing *CD74.* Besides that, we also found unique deletions of 6p21.2 containing *CDKN1A*, 11q12.3 containing *SCYL1*, 1q21.1 containing *NTRK1* and *MDM4,* and 6q21 containing *PRDM1.* It was found that the gain in 5p15.33 was one of the regions that reproducibly associates with lung cancer risk(Pande et al., 2011) and the gain in 20q13.33 was the main chromosomal abnormalities in colorectal carcinoma(Bui et al., 2020).

SCNA regions identified in this study encompassed genes that were known to driver metastasis. For instance, *CDKN2A* was frequently involved in genomic amplifications and deletions, respectively, as previously identified in a sequencing study of BrMs from patients with LUAD(Shih et al., 2020). *SCYL1* activates transcription of the telomerase reverse transcriptase and DNA polymerase beta genes. *SMAD4* is required for the function of *TGF-*β signaling pathway and related to carcinoma metastasis. *NTRK1* fusions trigger constitutive TRKA kinase activity(Vaishnavi et al., 2013), which activates cell growth and differentiation pathways(Alberti et al., 2003). *MDM4* is a p53 regulator and acts as an oncogene through p53-independent pathways(Gao et al., 2019). *PRDM1* gene encodes a protein that acts as a repressor of beta-interferon gene expression and regulator of *TP53* activity pathway(Boi et al., 2015).

Together, the identification of SCNAs further indicated that primary lung cancer and BrM lesions had similar genome-wide landscape of SCNAs, while BrM lesions contained unique chromosomal gains and losses that might potentially contribute to the development of BrMs.

### OXPHOS is enriched in LC-BrM

To identify differentially expressed genes between primary lung cancers and BrMs, we first applied ComBat to adjust for batch effects resulting from two independent cohorts with parametric empirical Bayes frameworks(Zhang et al., 2020). The returned expression matrix was corrected, leading to a single cohort of 56 pairs of primary lung cancers and BrM lesions (**Supplemental Fig. 5a and b**). Subsequently, we identified a total of 108 differentially expressed genes (DEGs) that were significantly up-regulated in BrM lesions (Log_2_Fold Change > 1 and adjusted Wald test p < 0.05) (**Fig. 5a**). To identify gene sets among the MSigDB Reactome collection which were positively correlated with the phenotype of BrM, we implemented gene set enrichment analysis (GSEA) which was based on the ranked gene list of DEGs. We identified 298 significantly enriched pathways (normalized enrichment score > 0 and adjusted p-value < 0.05), ranked the filtered pathways by normalized enrichment score (NES), and focused on the top 20 ranked pathways for the further analysis (**Supplemental Fig. 5c**). Among the top 20 ranked pathways, 5 of them were related to mitochondrial biogenesis and OXPHOS, including (1) *THE CITRIC ACID TCA CYCLE*, (2) *RESPIRATORY ELECTRON TRANSPORT*, (3) *RESPIRATORY ELECTRON TRANSPORT ATP SYNTHESIS BY CHEMIOSMOTIC COUPLING AND HEAT PRODUCTION BY UNCOUPLING PROTEINS*, (4) *COMPLEX I BIOGENESIS*, and (5) *PYRUVATE METABOLISM* (**Fig. 5b**).

**Figure 5.**
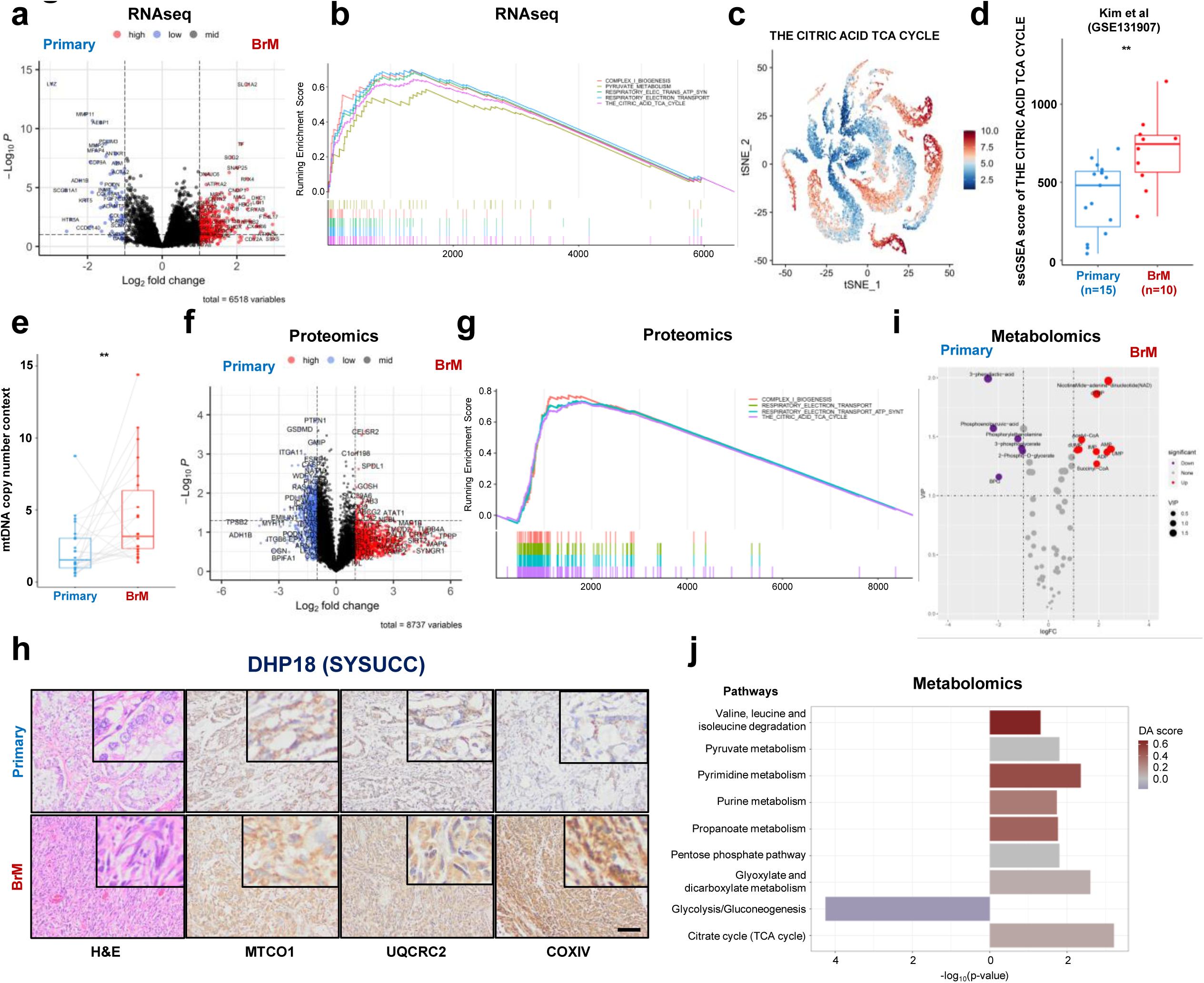
Oxidative phosphorylation was enriched in lung cancer brain metastases (LC-BrMs). **a**. The volcano plot of differentially expressed genes between primary lung cancer and LC-BrMs in two cohorts of 56 patients. **b**. Gene Set Enrichment Analysis (GSEA) plots of 5 mitochondrial pathways. The peak point of the top part in each plot represented the enrichment score (ES), whereas the bottom part showed where the rest of genes related to the pathway were located according to the ranking. **c and d**. The tSNE plot (**c**) and box plot (**d**) of enrichment score of the citric acid TCA cycle pathway in single cells of primary lung cancer and brain metastasis. Each dot represented an individual patient in panel d. The p-value was determined using two-sided Student’s t test. **e**. The box plot of relative mitochondrial DNA (mtDNA) content in paired primary lung cancers and brain metastasis lesions. The p-value was determined using pairwise two-sided Student’s t test. **f.** The volcano plot of differentially expressed proteins between primary lung cancer and LC-BrMs. **g**. GSEA plot of 4 mitochondrial pathways. The peak point of the top part in each plot represented the ES, whereas the bottom part showed where the rest of proteins related to the pathway were located according to the ranking. **h**. H&E and immunohistochemistry (IHC) staining of MTCO1, UQCRC2 and COXIV in a representative patient (DHP18) with paired primary lung cancer and LC-BrM lesions. **i.** The volcano plot of differentially expressed metabolites between primary lung cancer and LC-BrMs. **j.** Box plots for pathway analysis of differentially abundant metabolites between primary lung cancer and LC-BrMs.

Next, we validated these results by interrogating an independent cohort of scRNAseq(Kim et al., 2020) data that consisted of 45,149 single cells derived from 15 primary lung cancers and of 29,060 single cells derived from 10 BrM lesions (**Supplemental Fig. 5d**). We performed single sample gene set enrichment analysis (ssGSEA) and revealed that (1) all of these 5 mitochondrial pathways were indeed significantly enriched in BrM lesions as compared to primary lung cancers; and (2) ssGSEA scores of each of 5 mitochondrial pathways were significantly higher in BrM lesions compared to those of primary lung cancers (**Fig. 5c and d**, **Supplemental Fig 5e-l**). And this result was also observed in an independent lung cancer BrM cohort with spatial transcriptomic data available (Zhang et al., 2022) (**Supplemental Fig 5m**).

mtDNA plays a critical role in encoding many proteins that are important for the assembly, activity and function of mitochondrial respiratory complexes. To functionally validate whether mitochondrial biogenesis is enhanced in BrM lesions, we determined the relative mtDNA copy number in 24 pairs of primary lung cancers and BrM lesions derived from the SYSUCC cohort, for which genomic DNA were purified and available to us. Among 24 pairs, mtDNA copy number was significantly higher in 17 (70.8%) BrM lesions as compared to primary lung cancers (**Fig. 5e, Supplemental Fig 5n**).

To substantiate our findings on the mRNA level, we further performed the unbiased and global proteomics analysis using fresh frozen specimens of 4 pairs of primary lung cancers and BrM lesions. A total of 117 differentially expressed proteins were significantly up-regulated in BrM lesions (log_2_Fold Change > 1 and Wald test p < 0.05) (**Fig. 5f**). To identify pathways which were positively enriched in BrMs, we implemented the protein enriched analysis which was based on the ranked list of differentially expressed proteins. We identified 141 significantly up-regulated pathways (enrichment score > 0 and FDR < 0.05), ranked the filtered pathways by NES, and focused on the top 20 ranked pathways for further analysis (**Supplemental Fig. 5o**). Among the top 20 ranked pathways, 4 pathways were related to mitochondrial biogenesis and OXPHOS, including (1) *COMPLEX I BIOGENESIS*, (2) *RESPIRATORY ELECTRON TRANSPORT*, (3) *RESPIRATORY ELECTRON TRANSPORT ATP SYNTHESIS BY CHEMIOSMOTIC COUPLING AND HEAT PRODUCTION BY UNCOUPLING PROTEINS*, and (4) *THE CITRIC ACID TCA CYCLE* (**Fig. 5g**).

Additionally, we analyzed the expression of 3 representative respiratory chain complex related proteins, including MTCO1, UQCRC2 and COXIV, between primary lung cancer and BrMs by taking the advantage of RPPA data available from the Fukumura *et al*. cohort. Expression levels of MTCO1, UQCRC2 and COXIV were higher in BrM lesions as compared with primary lung cancers (**Supplemental Fig. 5p**). Moreover, we performed the IHC staining of 49 pairs of primary lung cancers and BrM lesions from both SYSUCC and WCH cohorts in order to independently validate the expression of MTCO1, UQCRC2 and COXIV at the protein level (**Fig. 5h**). As expected, we observed significantly higher IHC scores of MTCO1, UQCRC2 and COXIV in BrM lesions as compared to paired primary lung tumors (**Supplemental Fig. 5q**).

To further elucidate mitochondrial metabolism rewired in BrMs as compared to primary lung cancers, we performed LC-MS/MS based targeted metabolomics to measure 68 metabolites of energy metabolism.

A total of 10 metabolites were upregulated in BrM tumors when compared to primary lung cancer tumors (**Fig. 5i**). The pathway analysis based on differentially expressed metabolites indicated that citrate cycle (TCA cycle) and pyruvate metabolism pathways were significantly enriched in BrM tumors, whereas glycolysis/gluconeogenesis pathways were significantly enriched in primary lung cancer tumors (**Fig. 5j**).

Taken together, our results based on integrated analyses of RNAseq, proteomics and targeted metabolite data suggested that mitochondrial-specific metabolic adaptation was activated in BrMs of lung cancer.

### LC-BrMs present an immune suppressive microenvironment

We also identified a total of 119 DEGs that were significantly down-regulated in BrM tumors (Log_2_FoldChange < −1 and adjusted Wald test p < 0.05) (**Fig. 5a**). GSEA based on the ranked gene list of DEGs identified 126 pathways that were significantly down-regulated in BrM lesions (normalized enrichment score < 0 and adjusted p-value < 0.05). We also ranked the filtered pathways by NES and focused on the top 20 ranked pathways (**Supplemental Figure 6a**). Among the top 20 ranked pathways, 5 pathways that were significantly decreased in BrM tumors were related to signaling pathways of immune system, including (1) *INTERFERON ALPHA BETA SIGNALING*, (2) *INTERLEUKIN 2 FAMILY SIGNALING*, (3) *INTERFERON GAMMA SIGNALING*, (4) *IMMUNOREGULATORY INTERACTIONS LYMPHOID*, and (5) *INTERLEUKIN 4 AND 13 SIGNALING* (**Fig. 6a**).

**Figure 6.**
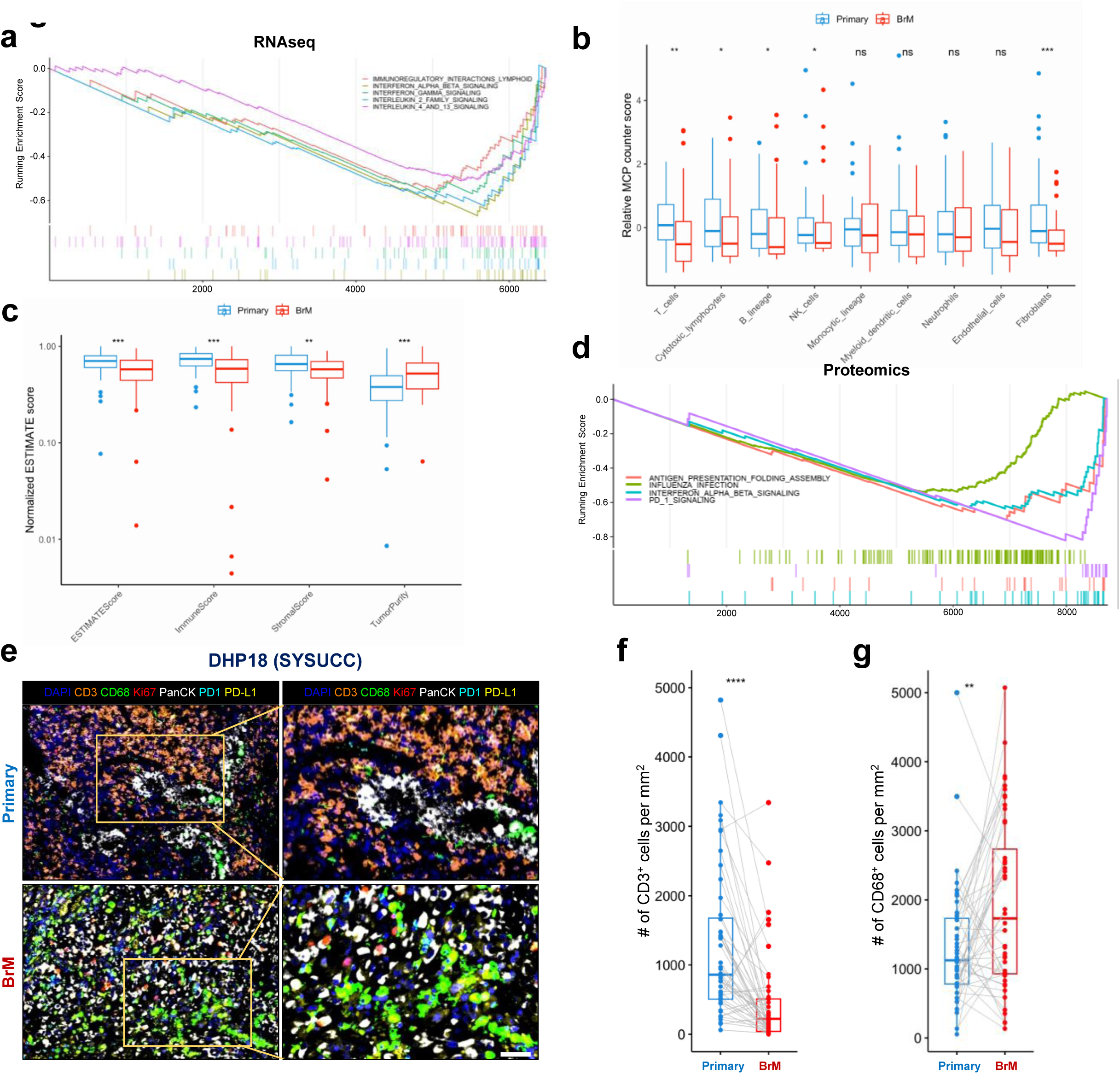
Brain metastasis (BrM) lesions presented an immunosuppressive tumor microenvironment. **a**. Gene Set Enrichment Analysis (GSEA) plots of 5 immune-related signaling pathways based on RNA sequencing data. The peak point of the top park in each plot represented the enrichment score (ES), whereas the bottom part showed where the rest of genes of each pathway were located according to the ranking. **b** and **c**. Box plot representation of normalized MCP counter scores (**b**) and ESTIMATE scores (**c**) for paired primary lung cancers and BrMs. The p*-*value was determined by the pairwise t-test. **d.** Protein set enriched analysis of 4 immune related signaling pathways based on proteomics data. The peak point of the top park in each plot represented the ES, whereas the bottom part showed where the rest of proteins of each pathway were located according to the ranking. **e**. Representative multiplex immunofluorescence (mIF) staining of paired primary lung cancer and BrM lesions from the patient DHP18. mIF markers include DAPI (blue), CD3 (orange), CD68 (green), Ki-67 (red), pan-cytokeratin (white), PD-1 (Cyan) and PD-L1 (yellow). Scale = 50[μm. **f** and **g**. Box plot representation of the count number of CD3^+^ (**f**) and CD68^+^ (**g**) cells per mm^2^ in paired primary lung cancer and BrM lesions (n=50 patients). The p-value was determined by the pairwise two-sided Student’s t test (*, p<0.05; **, p<0.01; ***, p<0.001; ****, p<0.0001).

To investigate the difference of immune infiltration between primary lung cancers and BrM lesions, we further performed computational analyses of RNAseq data using four algorithms, including (1) Estimation of Stromal and Immune cells in Malignant Tumor tissues using Expression data (ESTIMATE)(Yoshihara et al., 2013), (2) Microenvironment Cell Populations-counter (MCP-counter)(Becht et al., 2016), (3) a new version of Cell-type Identification By Estimating Relative Subsets Of RNA Transcripts (CIBERSORTx)(Newman et al., 2015; Newman et al., 2019), and (4) xCell(Aran et al., 2017), respectively. The result of MCP-counter demonstrated that BrM lesions exhibited significantly lower scores of T cells, cytotoxic lymphocytes, B lineage, NK cells, and fibroblasts as compared to primary lung tumors (**Fig. 6b**). The result of ESTIMATE indicated that BrM lesions showed significantly higher tumor purity scores but lower ESTIMATE scores, immune scores and stromal scores as compared to primary lung cancers (**Fig. 6c**). The result of xCell suggested that scores of several types of T cells, including CD4^+^ T cells and CD8^+^ T cells, were significantly lower in BrM tumors when compared to primary lung cancer lesions (**Supplemental Fig. 6b**). And the result of CIBERSORTx revealed that BrM lesions exhibited significantly higher scores of gamma delta T cells, but lower scores of monocytes, NK cells activated and CD8 T cells when compared to primary lung tumors (**Supplemental Fig. 6c**).

Similarly, we validated these results using the aforementioned independent scRNAseq dataset. Although ssGSEA scores of 4 out of 5 immune pathways were lower in BrM tumors compared to those of primary lung cancer tumors, we did not observe significant differences (**Supplemental Fig. 6d-n**).

The analysis of proteomics data from 4 pairs of fresh frozen primary lung cancers and BrM lesions also identified 756 differentially expressed proteins were significantly down-regulated in BrM lesions (log_2_Fold Change > 1 and Wald test p < 0.05) (**Fig. 5f**). We further performed protein enriched analysis based on the ranked list of differentially expressed proteins. We identified 247 significantly down-regulated pathways (enrichment score < 0 and FDR < 0.05), ranked the filtered pathways by NES, and focused on the top 20 ranked pathways for further analysis (**Supplemental Fig. 6o**). Among the top 20 ranked pathways, 4 pathways were related to immune regulation, including (1) *ANTIGEN PRESENTATION FOLDING ASSEMBLY*, (2) *INFLUENZA INFECTION*, (3) *INTERFERON ALPHA BETA SIGNALING*, and (4) *PD-1 SIGNALING* (**Fig. 6d**).

In order to experimentally validate results of computational analyses, we performed the mIF staining to unravel the immune microenvironment of BrM lesions. Towards that goal, we stained 48 pairs of primary and BrM tumors from both SYSUCC and WCH cohorts with CD3, CD68, PD-1, PD-L1, Pan-CK and Ki-67antibodies. As expected, BrM tumors exhibited significantly less infiltration of CD3^+^ immune cells but abundantly more CD68^+^ tumor-associated macrophages compared to primary lung cancer tumors (**Fig. 6e-g**).

Collectively, our integrated analyses of RNAseq, proteomics and mIF staining data suggested that BrM lesions were characterized by an immunosuppressive microenvironment.

### Gamitrinib exhibits its anti-tumor activity by inhibiting OXPHOS in BrM PDOs

Considering transcriptional alterations exhibited by BrM tumors as compared to paired primary lung cancers, we asked whether there was a phenomenon of transcriptional subtype switch unique to pairs of primary lung cancers and BrM lesions. Four tumor intrinsic (TI) subtypes of lung cancer were previously defined, including TI1 forlung dedifferentiated carcinoma, TI2 for lung adenocarcinoma, TI3 for lung squamous cell carcinoma, and TI4 for lung large cell neuroendocrine carcinoma(Ravi et al., 2023). Notably, the subtype switch from primary lung cancer to BrM occurred in 50% (28/56) patients (**Fig. 7a**).

**Figure 7.**
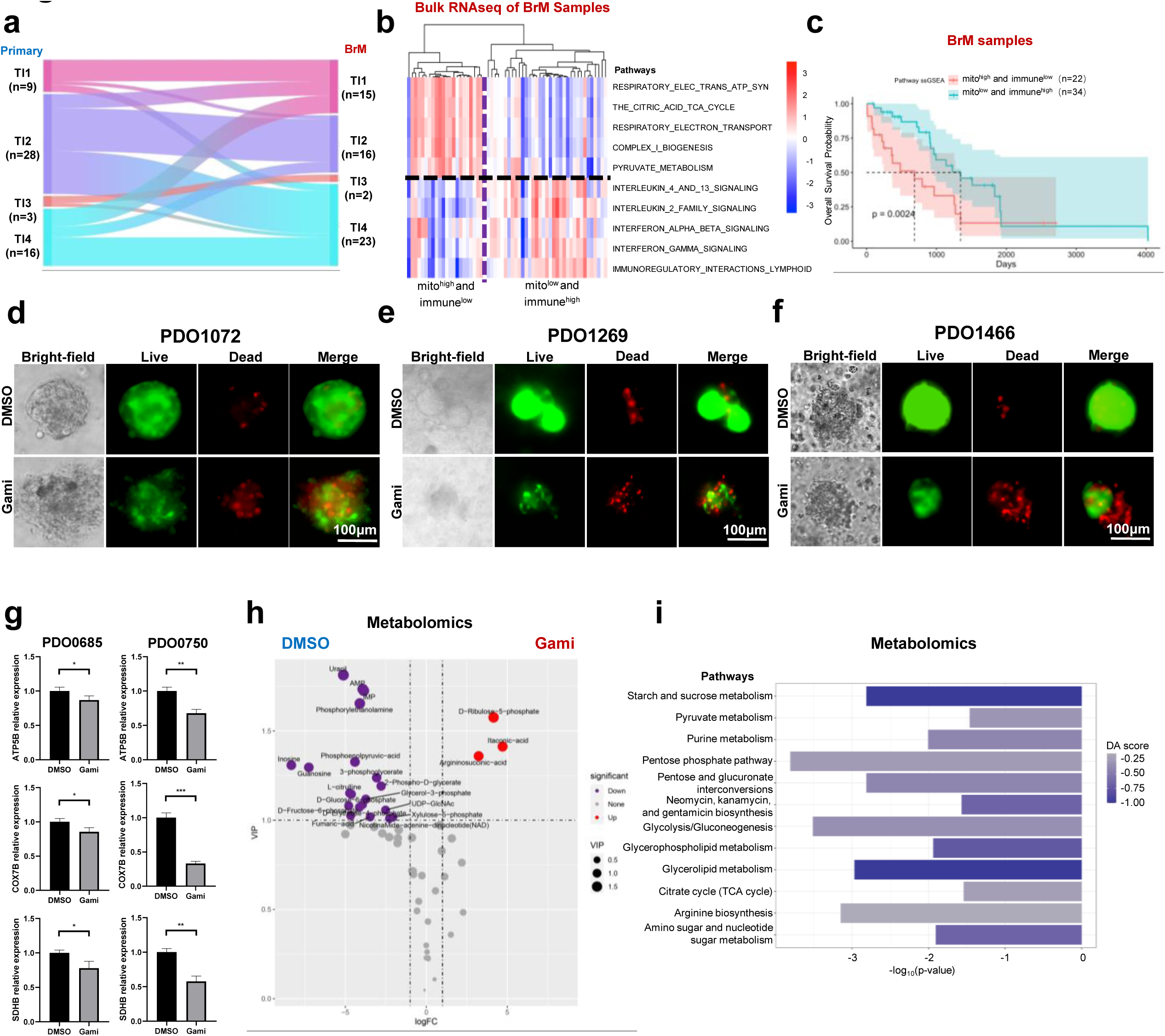
Gamitrinib exhibits its anti-tumor activity by inhibiting oxidative phosphorylation (OXPHOS) in patient-derived organoids (PDOs) of lung cancer brain metastasis (LC-BrM). **a.** Sankey plot indicating the transcriptional subtype switch from paired primary lung cancers to brain metastasis lesions. **b**. The heatmap showing the enrichment of 5 mitochondrial pathways and 5 immune pathways in LC-BrM lesions with RNA sequencing data. **c.** Kaplan-Meier survival plot of patients with LC-BrM lesions who were stratified into low and high subgroups based on single sample gene set enrichment analysis (ssGSEA) scores of mitochondrial pathways and immune pathways. The median ssGSEA score was used to define low and high subgroups. The p-value was computed by a two-sided log-rank test. **d-f.** Microscopic images of bright-field and immunofluorescence staining of PDO1072 (**d**), PDO1269 (**e**) and PDO1466 (**f**) PDOs which were treated with DMSO or gamitirnib. **g.** Relative mRNA expression of OXPHOS genes in two BrM PDOs (PDO0685 and PDO0750) treated with DMSO or gamitrinib. **h.** The volcano plot of differential metabolite analysis of LC-BrMs PDOs treated with DMSO or gamitrinib. **i.** Box plots for pathway analysis of differentially abundant metabolites of LC-BrMs treated with DMSO or gamitrinib.

Next, we calculated ssGSEA scores of 5 mitochondrial pathways and 5 immune pathways for each patient in Fukumura and SYSUCC cohorts. Interestingly, we observed there was an inverse correlation between immune and OXPHOS signaling pathways, suggesting that there was a subset of BrM lesions with activated mitochondrial biogenesis and OXPHOS but suppressed immune microenvironment (**Fig. 7b**). And this result was also observed in an independent cohort of 63 BrM lesions with mRNA microarray data available(Pocha et al., 2020) (**Supplemental Fig. 7a**). Next, we divided patients with LC-BrMs into two groups, mitochondria^low^/immune^high^ and mitochondria^high^/immune^low^ groups, according to ssGSEA scores, and performed Kaplan-Meier survival analysis. Indeed, patients in the mitochondria^low^/immune^high^ group had improved survival outcomes when compared to those in mitochondria^high^/immune^low^ group (**Fig. 7c**).

Since OXPHOS is enriched in LC-BrM tumors, we wondered whether OXPHOS inhibition would have anti-tumor activity in LC-BrMs. First, we tested the therapeutic efficacy of a specific OXPHOS inhibitor, gamitrinib(Chae et al., 2013; Kang et al., 2009; Wei et al., 2022; Zhang et al., 2016) in BrM PDOs, and the results indicated that all of the BrM PDOs were sensitive to gamitrinib with a median IC_50_ of 1.06 μM (**Supplemental Fig. 7b**). We further performed the live and dead staining using three representative BrM PDOs upon treatment with gamitrinib for 4 days. Our results suggested that gamitrinib induced cell death in PDOs (**Fig. 7d-f**). To investigate whether gamitrinib could inhibit OXPHOS in BrM PDOs, we performed RT-qPCR using PDOs treated with DMSO or gamitrinib. Indeed, the expression of 3 OXPHOS representative genes, including *ATP5B*, *COX7B* and *SDHB* were significantly decreased in BrM PDOs upon treatment with gamitrinib (**Fig. 7g**). Not surprisingly, the analysis of targeted metabolomics data further confirmed that OXPHOS-related pathways, such as TCA cycle and pyruvate metabolism were significantly inhibited in BrM PDOs upon treatment with gamitrinib (**Fig. 7h**). Taken together, our results demonstrated that gamitrinib could exhibit its anti-tumor activity by inhibiting OXPHOS in BrM PDOs.

### OXPHOS inhibition and anti-PD-1 treatment improve survival of mice with LC-BrMs

To identify actionable targets for LC-BrMs, we first generated an orthotopic mouse model of LC-BrM with Lewis lung cancer (LLC) cell line. LLC cells were injected via carotid artery and the *in vivo* establishment of LC-BrMs was further confirmed by bioluminescence imaging (BLI) (**Fig. 8a**). Since our integrated analyses established that mitchondrial biogenesis was activated and immune signaling pathways were suppressed in LC-BrMs, we sought to examine the efficacy of gamitrinib(Chae et al., 2013; Kang et al., 2009; Wei et al., 2022; Zhang et al., 2016) in combination with the immune checkpoint blockade anti-PD-1 antibody in targeting LC-BrMs. Toward that goal, mice were then randomly assigned to four treatment groups, including (1) control, (2) gamitrinib 10mg/kg, (3) anti-PD-1 10mg/kg and (4) the combination of gamitrinib 10mg/kg plus anti-PD-1 10mg/kg. As expected, gamitrinib as a monotherapy significantly improved survival of mice bearing LC-BrMs when compared to the control (log-rank test, p=0.01). Similarly, anti-PD-1 as a monotherapy also significantly delayed the death of mice bearing LC-BrMs (log-rank test, p=0.04). Remarkably, the rationale-based combination therapy of gamitrinib plus anti-PD-1 significantly prolonged survival of mice bearing LC-BrMs, leading to the longest median survival (log-rank test, p<0.001) (**Fig. 8b**). In comparison with mice treated by each monotherapy, those treated by the combination therapy of gamitrinib plus anti-PD-1 antibody had a trend of a longer survival; however, that survival difference did not reach the statistical significance (log-rank test, p<0.05).

**Figure 8.**
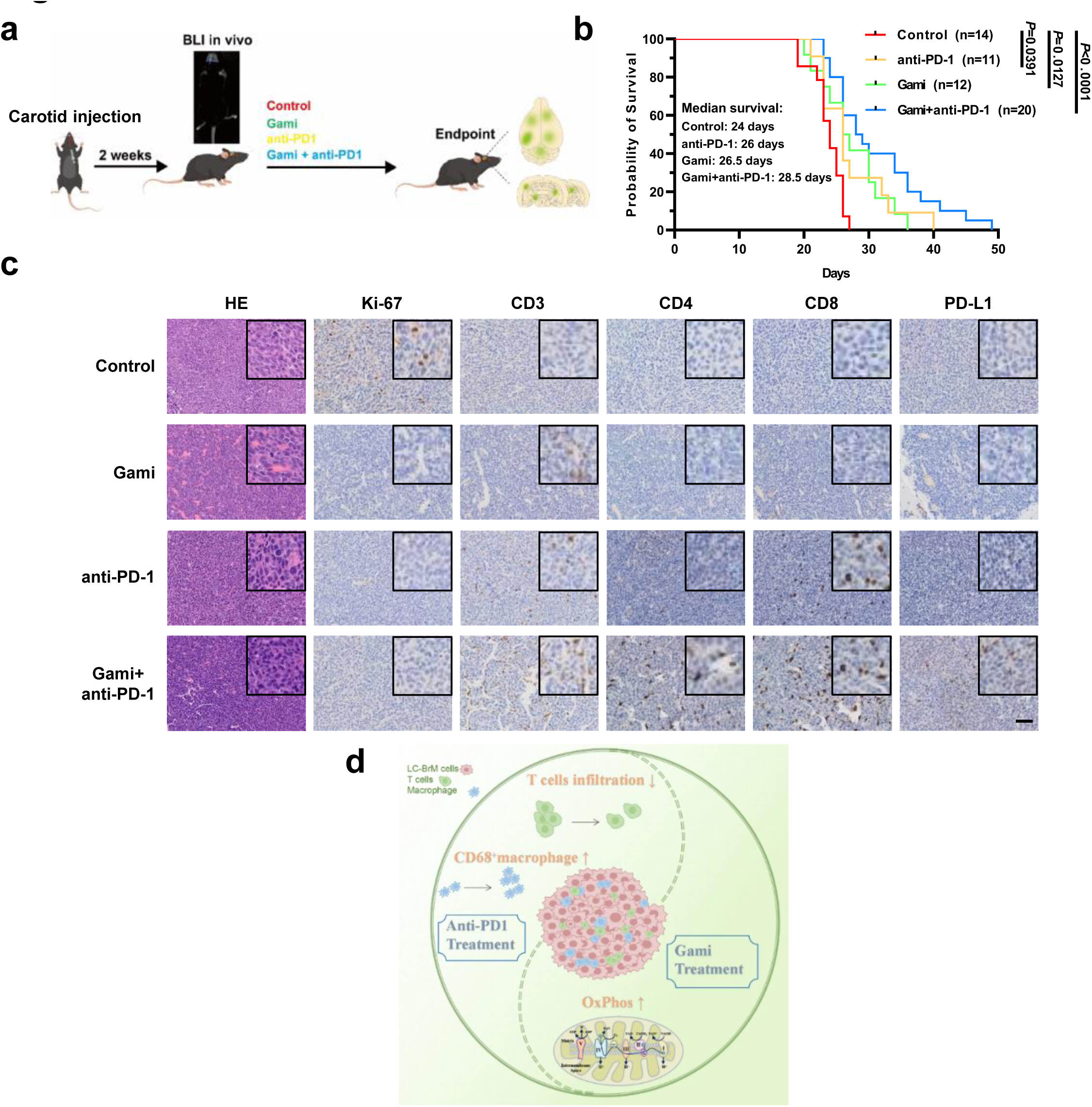
The combination therapy of an oxidative phosphorylation (OXPHOS) inhibitor plus anti-PD-1 blockade improved survival of mice with lung cancer brain metastases (LC-BrMs). **a.** Schematic illustration of the establishment of murine BrMs of Lewis lung cancer cells and treatment design. **b**. Kaplan-Meier survival plot of mice with LC-BrMs treated with control, gamitrinib at 10mg/kg, anti-PD-1 10mg/kg and gamitrinib 10mg/kg plus anti-PD-1 10mg/kg. The p*-*value is computed using a two-sided log-rank test. **c.** Representative images of H&E staining and immunohistochemistry (IHC) staining with anti-Ki-67, anti-CD3, anti-CD4, anti-CD8 and anti-PD-L1 for tumors in each treatment group. Scale = 50[μm. **d**. Schematic diagram of the current study and potential therapeutic implications.

Next, we analyzed representative tumor specimens from each treatment group with IHC staining. The IHC staining with Ki-67 antibody indicated that gamitrinib and/or anti-PD-1 inhibited tumor cell proliferation **(Fig. 8c, Supplemental Fig. 8a)**. IHC staining with CD3, CD4 and CD8 antibodies demonstrated that the infiltration of T cells in LC-BrMs was increased after treatment with anti-PD-1 antibody or the combination therapy of anti-PD-1 plus gamitrinib **(Fig. 8c, Supplemental Fig. 8b-d)**. Our results indicated anti-PD-1 antibody was able to enter the brain to enhance immune response. Moreover, expression of PD-L1 was upregulated in BrMs after treatment with gamitrinib or the combination therapy of gamitrinib plus anti-PD-1 antibody **(Fig. 8c, Supplemental Fig. 8e)**, further strengthening the rationale of combination therapy of gamitrinib plus anti-PD-1 antibody.

Taken together, these results indicated that the combination of OXPHOS inhibition and anti-PD-1 treatment is a promising therapeutic strategy for LC-BrMs (**Fig. 8d**).

## Discussion

Although previous studies have described the genomic and transcriptomic characteristics of LC-BrMs, molecular mechanisms underlying the biology of BrMs remain elusive. In this study, we performed comprehensive analyses of genomic, transcriptomic, proteomic and metabolomic data on both bulky tumor and single cell levels derived from four published and two newly generated patient cohorts, to get a global and deep understanding of molecular mechanisms and tumor microenvironment of LC-BrMs. We further validated multi-omics results by performing IHC and mIF stainings of patients’ tumor specimens, as well as in vitro and in vivo experiments of PDOs and mouse LC-BrM models. To our best knowledge, we assembled the largest cohort of paired primary lung cancers and BrMs with bulk multi-omic data available that permitted us to unravel the unique biology of BrMs derived from primary lung cancers. Our results reported significant genomic differences between primary lung cancers and BrMs, which surprisingly exhibited lower TMB and higher ITH in the latter. Additionally, we found that the mRNA expression signatures in primary tumors interconverted in the matched BrMs. Our study not only strengthened the fact that OXPHOS was elevated and immune activity was reduced in BrM but also presented a novel therapeutic intervention to delay the growth of BrM by the combination therapy of gamitrinib plus anti-PD-1 immunotherapy.

More than half of LC-BrMs harbor oncogenic driver mutations or CNVs. The amplifications of *MYC*, *YAP1* and *MMP13*, as well as the deletion of *CDKN2A/B* contributing to LC-BrMs originated from lung adenocarcinoma(Shih et al., 2020). In the current study, CNV analysis has identified unique amplification of *MDM4* and *NTRK1*, and deletion of *PRDM1, TNFAIP3, FANCC, TSC1* and *SMAD4* in LC-BrMs. Down-regulation of *SMAD4* was associated with the metastasis of pancreatic ductal adenocarcinoma (PDAC), and metastasis of *SMAD4*-negative PDAC was preferentially derived from the induction of mitochondrial OXPHOS (Liang et al., 2020).

Tumor heterogeneity poses severe challenges for cancer management. ITH representing the existence of genetic, epigenetic, and environmental heterogeneity within each individual tumor results in phenotypic heterogeneity and cellular plasticity, providing multiple mechanisms of therapeutic resistance and forming a highly adaptable and resilient disease. Our results showed that the ITH presented in LC-BrMs on the genetic level was higher than that in paired primary lung cancers, which was consistent with other studies. A study in which WES of primary lung–BrM paires was carried out showed that BrMs exhibited higher somatic variants and chromosomal alterations than primary lung cancers, particularly in genes associated with lung cancer (e.g., *KRAS, ROS1*, and *STK11*)(Tomasini et al., 2020). Liu *et al*. assessed the ITH of LC-BrMs by using scRNAseq data. Based on the differential gene expression analysis, tumor cells within the same LC-BrM sample could be clustered into 4 different subgroups which were enriched in genes related to OXPHOS, prostanoid biosynthesis and metabolic processes, and immune responses(Liu et al., 2020). Intriguingly, the high ITH is also a remarkable characteristic of gliomas which are one of the most common and deadly types of primary brain tumors(Nicholson and Fine, 2021). Thus, it is conceivable that the common brain-specific microenvironment shared by LC-BrMs and gliomas drives the co-evolution of cancers of different ontogenies and tumor microenvironment. Both LC-BrMs and gliomas are known to compromise the integrity of the BBB, resulting in a highly heterogeneous vasculature characterized by numerous distinct features, including non-uniform permeability and active efflux of molecules(Arvanitis et al., 2020). As the vasculature is dramatically changed during tumor growth and expansion, nutritional, oxygenic and metabolic properties are increasingly different in the tumor core, as compared to the periphery of the tumor and the neuroparenchyma harbouring an intact BBB(Arvanitis et al., 2020; Quail and Joyce, 2017). The ITH of microenvironment shaped by the heterogeneous vasculature may drive the selection of a diversified pool of clones that can successfully repopulate, resulting in genetically high ITH in LC-BrMs. An improved understanding of ITH in LC-BrMs will ultimately facilitate personalized medicine.

OXPHOS is the metabolic pathway in which cells use enzymes to oxidize nutrients, thereby releasing chemical energy in order to produce ATP. In eukaryotes, this takes place inside mitochondria. Previous studies observed that cancer cells universally down-regulate OXPHOS and up-regulate glycolysis compared with normal cells due to mutations in mtDNA or reduced mtDNA content in cancer cells(Gaude and Frezza, 2016; Larman et al., 2012; Viale et al., 2015; Yu, 2011). However, this assumption is being challenged by an increasing body of evidence that suggests that mitochondrial metabolism is not usually impaired, and OXPHOS can be also up-regulated in certain types of cancers under the stress stimuli or even in the face of active glycolysis(Birkenmeier et al., 2016; Vaupel and Mayer, 2012; Weinberg and Chandel, 2015; Whitaker-Menezes et al., 2011; Zacksenhaus et al., 2017). Moreover, metabolic heterogeneity has been demonstrated in tumors(Davidson et al., 2016; Hensley et al., 2016), and cancer stem cells with high metastatic and tumorigenic potential are more reliant upon OXPHOS than the bulk tumor populations(Viale et al., 2014). Recent studies demonstrated that significantly up-regulated OXPHOS was prominently featured in BrMs, and treatment with a direct OXPHOS inhibitor IACS-010759 as a monotherapy significantly hampered BrM formation in a murine model of breast cancer and prolonged survival of mice bearing melanoma BrMs(Fischer et al., 2019; Fukumura et al., 2021). Similarly, our results showed that LC-BrMs exhibited enhanced transcriptional signatures of OXPHOS and related biological processes involving the mitochondrial respiratory chain complex. The precise mechanisms driving up-regulation of OXPHOS in BrM are unclear and may vary by cancer type. Fukumura *et al*. found an increase in oxidative metabolism in brain metastatic derivatives of three distinct breast cancer cell lines, suggesting that up-regulation of OXPHOS in breast cancer cells is adaptively induced during the process of BrM(Fukumura et al., 2021). Melanoma BrMs appear to activate OXPHOS through up-regulation of the transcriptional regulator *PGC-1*α(Fischer et al., 2019). Fukumura *et al*. delineated links between the PI3K-AKT pathway and OXPHOS in breast, lung, and renal cell BrMs(Fukumura et al., 2021). In addition, the potential role of unique genetic mutations in BrMs in driving the up-regulation of OXPHOS is also worth to be further explored.

Tumor immune microenvironment is generally not concordant among the paired primary tumor and BrMs. In consistent with prior works(Fischer et al., 2019; Fukumura et al., 2021; Kim et al., 2020; Kudo et al., 2019), our analyses showed that the overall extent of infiltrating lymphocytes in BrMs is lower than that of the primary tumor, but that of macrophages is higher. Meanwhile, lower *PD-L1* expression in BrMs was discovered by us (data were not shown) and others(Mansfield et al., 2016). Lower infiltration of lymphocytes and expression of *PD-L1* in BrMs are associated with worse prognoses of patients and responses to immune checkpoint blockade(Berghoff et al., 2016; Goldberg et al., 2020; Zhou et al., 2018). These results have indicated the immune microenvironment of BrMs is overall suppressed compared to the primary tumor of lung. Discordances in the immune profile between BrMs and primary tumors can likely be explained by the significant selective pressure generated by the brain microenvironment, including the presence of the BBB and exclusive environmental cells(Quail and Joyce, 2017). In the normal brain, BBB remain as the initial gatekeeper of central nervous system (CNS) and is responsible for protecting CNS from a massive inflammation(Banks, 2016). Therefore, the healthy brain contains almost no lymphocytes, although there is evidence for immune surveillance of the normal human CNS by the lymphatic system in the brain(Loeffler et al., 2011; Louveau et al., 2015). As BrM lesions are established and further expand, the permeability of BBB is heterogeneously increased(Arvanitis et al., 2020). As a result, tumor-infiltrating lymphocytes and other blood-borne immune cells are observed in the BrMs but generally less than that observed in extracranial lesions. On the contrary, microglia are the resident macrophage cells located throughout the brain and spinal cord, which play a key role in overall brain maintenance. A large population of microglia-macrophages in brain maintenance usually exhibits a tumor-promoting phenotype, facilitating tumor progression, angiogenesis, immunosuppression, and therapeutic resistance(Mantovani et al., 2017; Qian and Pollard, 2010).

As solid tumors quickly proliferate and outgrow their chaotic vasculature, chronic hypoxia frequently occurs, which results from an imbalance between oxygen demand and poor oxygen supply due to abnormal vasculature(Dhani et al., 2015; Span and Bussink, 2015). Tumor hypoxia results in worse clinical outcomes because hypoxic areas are highly resistant to cancer therapy, including radiotherapy, targeted therapy and immunotherapy(Damgaci et al., 2018; Higgins et al., 2015; Overgaard, 2011). The absence of the oxygen enhancement effect mainly accounts for the resistance of hypoxic tumor cells to radiotherapy(Higgins et al., 2015). The immunosuppressive effect of hypoxia can result from both suppressed effects on immune effector cells and increased expression of cell-surface immune checkpoint molecules on tumor cells(Deng et al., 2019; Doedens et al., 2013; Iwai et al., 2002; Vito et al., 2020). Previous and present studies found that BrMs commonly up-regulated OXPHOS. OXPHOS inhibition could be an effective way to reduce the consumption of oxygen and to consequently reduce tumor hypoxia. Therefore, OXPHOS inhibition is emerging as an effective strategy to mitigate immune suppression and to enhance the efficacy of radio- and immuno-therapy in therapy-resistant hypoxic areas(Ashton et al., 2016; Chen et al., 2020; Chen et al., 2018; Najjar et al., 2019; Scharping et al., 2017; Zannella et al., 2013). Moreover, resistances to chemotherapy and targeted therapies appear to be generally coupled with an increase in OXPHOS, and OXPHOS inhibition overcomes resistance to docetaxel in prostate cancer, cytarabine in acute myeloid leukemia, 5-fluorouracil in colorectal and *MYC*/*PGC-1*α driven pancreatic cancer, to EGFR inhibition in *EGFR*-driven lung adenocarcinoma and MAPK or BRAF inhibition in *BRAF*-mutant melanoma(Adige et al., 2019; Bosc et al., 2017; Fischer et al., 2019; Vazquez et al., 2013; Yuan et al., 2013). Because tumors can display metabolic flexibility(Horsman et al., 2021; Martinez-Outschoorn et al., 2017), tumors with a high reliance on OXPHOS may be able to switch to glycolysis for ATP production. Strategies interfering with glycolysis may be employed to achieve synergistic combination effects with OXPHOS inhibitors(DeWaal et al., 2018; Sun et al., 2019). Above all, there is potential for combination of OXPHOS inhibitors with conventional chemotherapeutics, targeted therapies, radiotherapy, immunotherapy, and inhibitors of other metabolic pathways such as glycolysis for treating LC-BrMs.

There are some limitations of this study. Due to the retrospective nature, the clinicopathological information of some patients was unavailable. The correlation between mitochondrial and immune pathway scores with survival could be impacted by the treatment the patients received before surgical resection of BrMs. Given the heterogeneity of treatments prior to BrM resection and the small sample size of different treatment groups, we were unable to do analyses on the subgroup level. Additionally, we did not have patient-matched extracranial metastatic tissues to test the specificity of findings identified in BrMs. However, we believe these limitations will not compromise the reliability of our findings, which were explored by integrated analyses of multi-omics data and validated by PDOs and an orthotopic in vivo model of LC-BrM.

Given comprehensive analyses of the largest cohort of primary lung–brain metastasis pairs, and the novel and robust findings validated by multi-omic platforms, PDOs and a mouse LC-BrM model, our findings not only provide comprehensive and integrated perspectives of molecular underpinnings of LC-BrMs but also contribute to the development of a potential, rationale-based combinatorial therapeutic strategy of gamitrinib plus anti-PD-1 immunotherapy with the goal of translating it into clinical trials for patients with LC-BrMs.

## Methods

### Clinical samples

We included paired primary lung cancer and BrMs derived from three published patient datasets(Brastianos et al., 2015; Fukumura et al., 2021; Shih et al., 2020) and two newly generated cohorts in this study, in which multi-omics platforms were undertaken. Overall, we downloaded multi-omic data of primary lung–brain metastasis pairs from three eligible published cohorts, including Brastianos et al.(Brastianos et al., 2015), Shih et al.(Shih et al., 2020), and Fukumura et al.(Fukumura et al., 2021). Additionally, we generated two new cohorts derived from patients who were presented at SYSUCC and WCH. For the two newly generated cohorts, patient samples were collected under the Institutional Review Board (IRB) protocols of SYSUCC (Protocol B2021-256-01) and WCH (2019-57). Written informed consent were obtained from all patients. An additional published cohort of scRNAseq derived from unpaired primary lung cancers and BrMs was also included for validation analyses(Kim et al., 2020). No statistical methods were used to predetermine sample size. The experiments were not randomized, and the investigators were knowingly completing their work during experiments and outcome assessment.

Medical records and archived formalin fixed paraffin-embedded (FFPE) tissues of patients from SYSUCC and WCH were retrospectively retrieved. Clinicopathological data including patient age at diagnosis of BrMs, sex, smoking history, date of primary diagnosis, pathological diagnosis, cancer stage of primary diagnosis, date of BrMs diagnosis, treatment before BrMs diagnosis, treatment for BrMs before surgical resection, date of craniotomy, location of BrMs, date of last follow-up, deceased date, and survival status were collected from medical records. The last date of follow-up was December 2022. Survival status of patients was determined from clinical attendance records or direct telecommunication with patients or their families. OS was defined as the time between craniotomy for BrMs and cancer-caused death or the date of last follow-up.

For preparation of FFPE tumor sections of patients from SYSUCC and WCH, an experienced pathologist reviewed the H&E staining of the surgically resected tumor samples and selected the most representative tumor tissue of each sample. Then, the representative FFPE samples of which sizes were comparable were sectioned. Fresh-frozen whole blood samples of patients were obtained retrospectively from the biobank of SYSUCC.

### DNA extraction and WES

After FFPE sample sections were scalpeled into 1.5 mL micro centrifuge tube. Deparaffinization solution was used to remove paraffin. Then Maxwell 16 FFPE Plus LEV DNA Purification Kit (Promega) was used to extract FFPE DNA, and genomic DNA was extracted from a 0.5 mL aliquot of whole blood with the DNeasy Blood and Tissue Kit (69506, Qiagen) according the protocal’s instructions. Then the integrity and concentration of the total DNA was determined by agarose electrophoresis and Qubit 3.0 fluorometer dsDNA HS Assay (Thermo Fisher Scientific). About 300 ng high-quality DNA sample was used to construct sequencing library. The 300ng genomic DNA concentrations were sheared with Covaris LE220 Sonicator (Covaris) to target of 150-200bp average size. DNA libraries were prepared using SureselectXT reagent kit (Agilent). The fragments were repaired the 3’and 5’overhangs using End repair mix (component of SureselectXT) and purified using Agencourt AMPure XP Beads (Beckman). The purified fragments were added with ‘A’ tail using A tailing Mix (component of SureSelectXT) and then ligated with adapter using the DNA ligase (component of SureselectXT). The adapter-ligated DNA fragments were amplified with Herculase II Fusion DNA Polymerase (Agilent). Finally, the pre-capture libraries containing exome sequences were captured using SureSelect Human All Exon V6 kit (Agilent). DNA concentration of the enriched sequencing libraries was measured with the Qubit 3.0 fluorometer dsDNA HS Assay (Thermo Fisher Scientific). Size distribution of the resulting sequencing libraries was analyzed using Agilent BioAnalyzer 4200 (Agilent). Sequencing was performed using an NovaSeq 6000 S4 following Illumina-provided protocols for 2×150 paired-end sequencing in Mingma Technologies (Shanghai, China).

### RNA extraction and RNAseq

After FFPE sample sections were scalpeled into 1.5 mL micro centrifuge tube. Deparaffinization solution was used to remove paraffin. Then Maxwell 16 LEV RNA FFPE kit (Promega) was used to extract FFPE RNA according the protocal’s instructions. RNA integrity was determined by 2100/2200 Bioanalyser (Agilent) with DV200 (Percentage of RNA fragments > 200 nt fragment distribution value) and quantified using the NanoDrop (Thermo Scientific). RNA purification, reverse transcription, library construction and sequencing were performed at Mingma Technologies (Shanghai, China) according to the manufacturer’s instructions (Illumina). The captured coding regions of the transcriptome from total RNA were prepared using TruSeq® RNA Exome Library preparation Kit. For FFPE sample, RNA input for library construction was determined by the quality of RNA. Generally, 20 ng RNA was recommended for FFPE RNA sample with high quality and 20-40 ng RNA is for FFPE RNA sample with medium quality. For FFPE RNA sample with low quality, 40-100 ng total RNA was used as input. Then the cDNA was generated from the input RNA fragments using random priming during first and second strand synthesis and sequencing adapters were ligated to the resulting double-stranded cDNA fragments. The coding regions of the transcriptome were then captured from this library using sequence-specific probes to create the final library. After library constructed, Qubit 2.0 fluorometer dsDNA HS Assay (Thermo Fisher Scientific) was used to quantify concentration of the resulting sequencing libraries, while the size distribution was analyzed using Agilent BioAnalyzer 2100 (Agilent). Sequencing was performed using an NovaSeq 6000 S4 following Illumina-provided protocols for 2×150 paired-end sequencing in Mingma Technologies at Shanghai, China.

### Computational pipelines

All pipelines were developed according to National Cancer Institute sequencing pipelines. Unless otherwise stated, all tools mentioned are part of GATK 4 suite. All data were analyzed with homogenous pipelines capable of processing raw fastq files as well as re-processing previously analyzed bam files.

### Alignment and pre-processing

WES data pre-processing was conducted in accordance to the GATK Best Practices using GATK 4.0. In brief, aligned BAM files were separated by read group, sanitized and stripped of alignments and attributes using ‘RevertSam’, which generated one unaligned BAM (uBAM) file per readgroup. Uniform readgroups were assigned to uBAM files using ‘AddOrReplaceReadgroups’. Then uBAM files were reverted to interleaved fastq format using ‘SamToFastq’. Unaligned fastq files underwent quality control using ‘FastQC’. Sequencing adapters were marked and removed using ‘Trim_glore. Fastq files were finally aligned to the b37 genome using ‘BWA MEM’ and attributes were restored using ‘MergeBamAlignment’. ‘MarkDuplicates’ was then used to merge aligned BAM files from multiple readgroups and to mark PCR and optical duplicates across identical sequencing libraries. Lastly, base recalibration was performed using ‘BaseRecaliBrator’ followed by ‘ApllyBQSR’. Coverage statistics were gathered using ‘CollectHsMetrics’. Quality control of alignment was performed by running ‘ValidateSamFile’ on the final BAM file and quality control results were further inspected using ‘MultiQC’. The tool ‘CrosscheckFingerprints’ was used to confirm that all readgroups within a sample belong to the same individual, and that all samples from one individual match. Any mismatches were marked and excluded from further analysis. RNA data pre-processing was conducted in accordance to the mRNA analysis pipeline published on National Cancer Institute. The raw fastq files went through quality control using ‘FastQC’. The bases that did not pass it were cut off using ‘Trim_glore’. Fastq files were then aligned to the b37 genome using ‘STAR’ to generate BAM files. Post quality control were applied to these BAM files using ‘CollectRNASeqMetrics’, which produced metrics describing the distribution of the bases within the transcripts.

### WES Variant detection

Variant detection was performed in accordance to the GATK Best practices using GATK4. Germline variants were called from control samples using Mutect2 in artefact detection mode and pooled into a cohort-wide panel of normal samples. Somatic variants were subsequently called from tumor samples with match control samples (tumor with matched normal mode) using Mutect2. The parameters in Mutect2 include matched normal sample, the reference fasta file, the panel of normal mentioned in the above and the gnomAD germline resources as additional controls. Cross-sample contamination was evaluated using ‘GetPileupSummaries’ and ‘CalculateContamination’ run for all samples, both tumour and matching normal. Read orientation artefacts were evaluated using ‘Collect-F1R2Counts’ and ‘LearnReadOrientationModel’. Additional filters were added through ‘FilterMutectCalls’, including artifact-in-normal and contamination fractions.

### WES variant post-processing

BCFTools was used to normalize, sort and index variants. A consensus VCF was generated from all variants in the cohort with any duplicate variants removed. The VCF file was annotated using GATK4.1 Funcotator and the v1.7.20200521s annotation data source.

### Mutational burden

The mutational burden was calculated as the number of mutations per Mb sequenced. A minimum coverage threshold of 15x was required for each base. tcgaCompare was used to compare BrM and primary against 33 TCGA cohorts including LUAD, LUSC, SKCM and so on.

### Unique and shared mutations

Post-processed mutations in tumor samples were compared with its matched ones from primary lung cancer samples. In between two mutation results, the shared mutations were defined as same mutations on same chromosome position and leading to same type variants on same genes. Otherwise, the rest variants were defined as unique to their own.

### Mutational signatures and Oncoplots

The relative contributions of the COSMIC mutational signatures were determined from mutations identified in a patient’s LC-BrM and primary lung cancer samples. Adjacent bases surrounding the mutated base was obtained and formed a mutation matrix. The matrix was used to run NMF and measures the goodness of fit, in terms of Cophenetic correlation. Then, the matrix was decomposed into multiple signatures, and compared to known signatures from COSMIC database depending on the calculated cosine similarity. All BrM samples and primary samples were compared to identify differentially mutated genes. First all called somatic variants were merged using ‘merge-vcf’, converted to one VCF file and further annotated using Funcotator. The differentially mutated genes were then detected using fisher test on all genes between two cohorts, and plotted using oncoplots.

### Tumor heterogeneity and MATH

The heterogeneity was inferred by clustering VAF in both LC-BrM and primary lung cancer samples. The median absolute deviation (MAD) was determined through mutant-allele fraction (MAF) for all tumors by calculating the absolute value of the difference of each MAF from the median MAF value. MATH score is a simple quantitative measure of ITH, which is the width of the VAF distribution and calculated as the percentage ratio to the MAD to the median of the distribution of MAFs among the tumor’s mutated genomic loci.

### Copy number segmentation

Copy number identification was performed according to recommended workflow (http://varscan.sourceforge.net/copy-number-calling.html) for the variant detection in massively parallel sequencing data (VarScan). Both LC-BrM and primary lung cancer samples went through the same workflow in order for us to identify tumor-specific (somatic) copy number changes. The raw copy number calls were determined by using ‘samtools mpileup’ on normal blood samples and tumor samples (both BrM and primary lung cancer). The GC content of raw copy number calls was adjusted and preliminary calls were made using ‘copyCaller’. Then circular binary segmentation algorithm was applied to adjusted copy number using DNAcopy library from BioConductor and the results was visualized using DNAcopy package. Finally, the data points were re-centered using ‘copyCaller’ again if the the segments were above or below the neutral value.

### Copy number calling

Copy number calling was performed using GISTIC2.0 to identify genes of SCNAs. Segmented copy-number (from Conflict-based search algorithm) was deconstructed into its most likely set of underlying SCNAs using ‘Ziggurat Deconstruction’ algorithm, which separates arm-level and focal SCNAs explicitly by length. Then the deletion and amplification scores for both focal and arm-level SCNAs were calculated through GISTIC probabilistic framework based on markers. Finally, the number of independently significant SCNAs on each chromosome was determined using the ‘Arbitrated Peel-off’ algorithm, and the boundaries of significantly altered regions were determined using ‘RegBounder’ approach based on approximating the amount of expected local variation in GISTIC score profiles.

We utilize two additional outputs, namely “Amp_genes.conf_90.txt” and “Del_genes.conf_90.txt,” to identify distinctive amplification and deletion peaks in BrM in comparison to the Primary. These outputs comprise a tabular presentation of amplification and deletion peaks, accompanied by the corresponding genes and their respective q-values. We have emphasized the peaks exclusively detected in BrM, with a q-value below 0.05.

To refine the selection of functional gene-level CNVs within the arm-level regions, we exclusively retained CNVs that directly impact the functionality of specific genes, encompassing both oncogenes and tumor suppressors. Moreover, we included CNVs labeled as oncogenic or predicted to be oncogenic according to OncoKB [29625050].

### Batch effect removal

RNAseq from three batches (Fukumura *et al*. batch, SYSUCC batch 1, and SYSUCC batch 2) was batch corrected using ComBat-seq, using a negative binomial regression model that retains the integer nature of count data in RNAseq. SYSUCC has two batches with 4 overlapped patients, so a Pearson correlation value was calculated for each of these 4 patients’ RNAseq counts in-between two batches, and all values are higher than 0.97. The RNAseq data from SYSUCC batch 2 was retained. Finally, the corrected counts’ data was plotted using PCA based on their similarities.

### Differentially expressed genes in SCNAs

Similar approach as stated in ‘Comparing two cohorts’ was used to detect the differentially expressed genes between BrM and primary lung SCNAs. The only difference is to replace the total number of somatic mutations to the total number of amplification and deletion on each gene.

### Differential gene expression analysis of RNAseq data

Following alignment, BAM files were processed through the RNA Expression Workflow to determine RNA expression levels. The reads mapped to each gene are enumerated using ‘HT-Seq-Count’. The number of reads mapped to each gene are normalized using ‘DESeq2’, which uses the negative binomial as the reference distribution and provides its own algorithm. The results from DESeq2 include base means across samples, log2 fold changes, standard errors, test statistics, p-valued and adjusted p-values. Visualization of these significant genes are plotted using ‘Volcano Plot’.

### Gene set enrichment analysis

Pathway analysis was conducted using ‘fgsea’, a fast preranked GSEA. The ranked significant genes collected from ‘DESeq2’ and reactome pathway dataset (c2.cp.reactome.v7.4) were given as inputs to ‘fgsea’, which generated outputs including pathway names, enrichment score, normalized enrichment score and its p-value.

### Immune cell abundance analysis

Relative immune cell fraction data used in downstream neoantigen analysis were determined by ‘MCPcounter’ R package. ESTIMATE relative immune cell analysis were determined by ‘Estimate’ r package. Gene expression data was used in CIBERSORTx to provide an estimation of the abundances of member cell types in a mixed cell population(https://cibersortx.stanford.edu/).

### Survival analysis

The OS defined as the time between craniotomy for BrMs and cancer-caused death or the date of last follow-up was subjected to survival analyses carried out using the Survminer package in R software (version 4.1.0.). Gehan-Breslow (a generalized Wilcoxon) tests were used for univariate comparisons in the Kaplan-Meier survival curve. The variate is the average of pathway enrichment scores that was calculated using method single sample GSEA in the Gene Set Variation Analysis package. Primary tumor or BrM samples were divided into two groups according to the scores, including enriched (greater than zero) and non-enriched (less than zero).

### Dimension reduction and unsupervised clustering for scRNAseq data

scRNAseq data was collected from a published dataset Kim 2020. It contained 208,506 single cells from LUAD patients, in which 29,060 were LC-BrM cells and 45,149 were primary lung cancer cells. Specifically, only the BrM and primary lung cancer cells were included for the downstream analysis. The data was downloaded as normalized log2TPM matrix, and the genes that were expressed at low levels were removed. Variably expressed genes with mean expression between 0.0125 and 3 were selected using ‘Seurat’ in R, and then used to compute the principal components (PCs). The significant PCs were selected using ‘PCElbowPlot’ and ‘JackStraw’ in Seurat. Cell clustering and tSNE visualization were performed using ‘FindClusters’ and ‘RunTSNE’ functions, respectively. Gene set enrichment was calculated using the ‘enrichIt’ function from R package ‘escape’ and displayed using ‘FeaturePlot’ with tSNE reduction.

### mtDNA copy number detection

20 ng FFPE genomic DNA was used to conduct RT-qPCR. mtDNA were amplified using specific primers with the Fast SYBR™ Green Master Mix (ThermoFisher, 4385610). Primers sequence for RT-qPCR were as follows: mtDNA: forward primer, 5’-CACCCAAGAACAGGGTTTGT-3’, and reverse primer, 5’-TGGCCATGGGTATGTTGTTA-3’; and β2-microglobulin: forward primer, 5’-TGCTGTCTCCATGTTTGATGTATCT-3’, and reverse primer, 5’-TCTCTGCTCCCCACCTCTAAGT-3’. The relative mtDNA copy number was analyzed by the 2^-ΔΔCt^ method.

### IHC staining

The FFPE sections were rewarmed at 65°C for 3 hours and then deparaffinized and rehydrated with degraded alcohol. After that, heat-induced antigen retrieval was carried out with 0.01 M citrate salt buffer (ZSGB-BIO, Beijing, China) at 95 °C for 15 minutes. After being incubated with 0.3% H_2_O_2_ for 10 min and blocked with 10% fetal calf serum for 15 minutes, the tissue sections were incubated with anti-MTCO1 (abcam, ab14705), anti-UQCRC2 (Proteintech, 14742-1), anti-COXIV (CST, 4850), anti-Ki-67 (abcam, ab16667), anti-CD3 (CST, 78588), anti-CD4 (abcam, ab183684), anti-CD8 (CST, 98941) and anti-PD-L1 (CST, 64988) antibodies at 4 °C overnight. Subsequently, these tissue sections were incubated with horseradish peroxidase-conjugated anti-mouse or anti-rabbit antibody (ZSGB-BIO, PV-6000) at room temperature for 60 minutes. Then, the sections were stained with DAB + substrate-chromogen solution (ZSGB-BIO, PV-6000) at room temperature for 30 seconds and counterstained with hematoxylin. The expression level of MTCO1, UQCRC2 and COXIV were evaluated by both staining intensity and percentage of staining positive cells according to a semi-quantitative scoring system. Staining intensity was scored as 0 for negative staining, 1 for weak staining, 2 for moderate staining, and 3 for strong staining. Percentage of positive cells was quantified as 0 for ≤5% positive cells, 1 for 6-25%, 2 for 26-50%, 3 for 51-75% and 4 for ≥76%. The immunoreactivity score was then generated by multiplying score of staining intensity and percentage of positive cells.

### mIF

Twenty-eight matched primary lung tumors and BrM lesions were stained with mIF. The formalin fixed paraffin embedded bullae were sectioned and processed using Opal Polaris™ 7-color Manual IHC Kit (Akoya Biosciences) following the manufacturer’s recommendation. The mIF panel included DAPI (Abcam, ab104139), anti-CD3 (Abcam, ab16669), anti-CD68 (Abcam, ab192847), anti-Ki-67 (Abcam, ab16667), anti-panCK (Abcam, ab7753), anti-PD-1 (Abcam, ab237728) and anti-PD-L1 (Abcam, ab237726).

### Proteomics assay

Fresh-frozen tumor samples were first grinded by liquid nitrogen and then the powder was transferred to a 1.5 ml centrifuge tube and sonicated three times on ice, using a high intensity ultrasonic processor in a lysis buffer (8M urea including 1mM PMSF and 2mM EDTA). Then, the remaining debris was removed by centrifugation at 15000g at 4°C for 10 min. Finally, the protein concentration was determined with a BCA kit according to the instructions of the manufacturer.

Equal amount of proteins from each sample were used for tryptic digestion. Add 8M urea to 200ul to the supernatants, then reduced with 10 mM DTT for 45 minutes at 37°C and alkylated with 50 mM iodoacetamide for 15 minutes in a dark room at room temperature. 4 × volume of chilled acetone was added and precipitated at −20°C for 2 hours. After centrifugation, the protein precipitate was air-dried and resuspended in 200ul of 25mM ammonium bicarbonate solution and 3ul of trypsin (Promega) and digested overnight at 37°C. After digestion, peptides were desalted using C18 Cartridge followed by drying with Vacuum concentration meter, concentrated by vacuum centrifugation and redissolved in 0.1% (v/V) formic acid.

Liquid chromatography (LC) was performed on a nanoElute UHPLC (Bruker Daltonics, Germany). About 200 ng peptides were separated within 60 min at a flow rate of 0.3 uL/min on a commercially available reverse-phase C18 column with an integrated CaptiveSpray Emitter (25 cm x 75 μm ID, 1.6 μ m, Aurora Series with CSI, IonOpticks, Australia). The separation temperature was kept by an integrated Toaster column oven at 50°C. Mobile phases A and B were produced with 0.1 vol.-% formic acid in water and 0.1% formic acid in HPLC-grade acetonitrile (ACN). Mobile phase B was increased from 2 to 22% over the first 45 min, increased to 35% over the next 5 min, further increased to 80% over the next 5 min, and then held at 80% for 5 min. The LC was coupled online to a hybrid timsTOF Pro2 (Bruker Daltonics, Germany) via a CaptiveSpray nano-electrospray ion source (CSI). The timsTOF Pro2 was operated in Data-Dependent Parallel Accumulation-Serial Fragmentation (PASEF) mode with 10 PASEF MS/MS frames in 1 complete frame. The capillary voltage was set to 1400 V, and the MS and MS/MS spectra were acquired from 100 to 1700 m/z. As for ion mobility range (1/K_0_), 0.7 to 1.4 Vs/Cm^2^ was used.

The TIMS accumulation and ramp time were both set to 100ms, which enable an operation at duty cycles close to 100%. The “target value” of 10,000 was applied to a repeated schedule, and the intensity threshold was set at 2500. The collision energy was ramped linearly as a function of mobility from 59 eV at 1/K_0_ = 1.6 Vs/Cm^2^ to 20 eV at 1/K_0_ = 0.6 Vs/Cm^2^. The quadrupole isolation width was set to 2Th for m/z < 700 and 3Th for m/z > 800.

MS raw data were analyzed using DIA-NN (v1.8.1) with library-free method. the Homo sapiens SwissProt database (20425 entries) was uesed to creat a spectra library with deep learning algrithms of neural networks. the option of MBR was employed to create a spectral library from DIA data and then reanlyse using this library. FDR of search results was adjusted to < 1% at both protein and precursor ion levels, the remaining identifications were used for further quantification analysis.

### Quantitative analysis of energy metabolism

ACN and methanol (MeOH) were purchased from Merck (Darmstadt, Germany). MilliQ water (Millipore, Bradford, USA) was used in all experiments. All of the standards were purchased from Sigma-Aldrich (St. Louis, MO, USA) and Zhenzhun etc. Formic acid was bought from Sigma-Aldrich (St. Louis, MO, USA). The stock solutions of standards were prepared at the concentration of 1 mg/mL in MeOH and other solutions. All stock solutions were stored at −20°C. The stock solutions were diluted with MeOH to working solutions before analysis.

After the fresh-frozen samples were thawed and smashed, an amount of 0.05 g of each sample was mixed with 500 μL of 70% methanol/water. The sample was vortexed for 3 min under the condition of 2500 r/min and centrifuged at 12000 r/min for 10 min at 4°C.Take 300 μL of supernatant into a new centrifuge tube and place the supernatant in −20°C refrigerator for 30 min, Then the supernatant was centrifuged again at 12000 r/min for 10 min at 4°C. After centrifugation, transfer 200 μL of supernatant through Protein Precipitation Plate for further LC-MS analysis.

The sample extracts were analyzed using an LC-ESI-MS/MS system (Waters ACQUITY H-Class; MS, QTRAP® 6500+ System). The analytical conditions were as follows. Amide method: HPLC: column, ACQUITY UPLC BEH Amide (i.d.2.1×100 mm, 1.7 μm); solvent system, water with 10mM Ammonium acetate and 0.3% Ammonium hydroxide (A), 90% acetonitrile/water (V/V)(B); The gradient was started at 95% B (0-1.2 min), decreased to 70% B (8 min),50% B (9-11 min), finally ramped back to 95% B (11.1-15 min); flow rate, 0.4 mL/min; temperature, 40°C; injection volume: 2 μL.

Linear ion trap and triple quadrupole scans were acquired on a triple quadrupole-linear ion trap mass spectrometer (QTRAP), QTRAP® 6500+ LC-MS/MS System, equipped with an ESI Turbo Ion-Spray interface, operating in both positive and negative ion mode and controlled by Analyst 1.6.3 software (Sciex). The ESI source operation parameters were as follows: ion source, ESI+/-; source temperature 550 ∘C; ion spray voltage (IS) 5500 V (Positive), −4500 V (Negative); curtain gas was set at 35 psi, respectively. Tryptophan and its metabolites were analyzed using scheduled multiple reaction monitoring (MRM). Data acquisitions were performed using Analyst 1.6.3 software (Sciex). Multiquant 3.0.3 software (Sciex) was used to quantify all metabolites. Mass spectrometer parameters including the declustering potentials (DP) and collision energies (CE) for individual MRM transitions were done with further DP and CE optimization. A specific set of MRM transitions were monitored for each period according to the metabolites eluted within this period.

### PDOs generation and viability assay

The culture of PDOs was performed according to the method previously reported(Wang et al., 2023). Briefly, freshly resected LC-BrMs were finely minced and transferred to a 50 ml conical tube, including a digestion mix 400 consisting of serum-free advanced DMEM/F-12 medium (Gibco, USA) and 1 mg/ml collagenase IV (Sigma, USA), and incubated for 1 h at 37 °C with shaking. The cell suspension that had been digested was combined with Matrigel (BD Biosciences, USA) at a ratio of 1:1.5 (v/v) and then placed in 96-well plates at a volume of 10μl per well. The culture medium contained advanced DMEM/F-12 with PS (1×), glutamine (1×), B27 supplement (1×), nicotinamide (5 mM), nacetylcysteine (1.25 mM), A83-01 (500 nM), SB202190 (500 nM), Y-27632 (5 mM), noggin (100 ng/ml), R-spondin 1 (250 ng/ml), FGF 2 (5 ng/ml), FGF 10 (10 ng/ml) and EGF (5 ng/ml). Supplemented culture medium was added 100ul to each per well, and organoids were maintained in a 37 °C humidified atmosphere under 5% CO2.

To measure the IC50 of gamitrinib on PDOs, PDOs were dissociated into smaller clusters containing approximately 2000 cells, resuspended in 36 µL culture medium and seeded in each well of a 384-well plate. After 48 h, 4 µL of a threefold dilution series of each drug was dispensed separately; three technical replicates of each drug were tested on three plates. After 3 days, cell viability was quantitated using the CellTiter-Glo 3D Cell Viability Assay (G9681, Promega) following the manufacturer’s instructions. Relative luminescence units (RLU) for each well were normalized to the median RLU from the DMSO control wells, used as 100% viability. IC50 values were generated using Prism 9 (GraphPad Software, Boston, MA, USA).

PDOs were seeded in 96-well plates with 5 μl of Matrigel per well in a total volume of 100 μl of the culture medium, and treated with gamitrinib at IC50 dosage or DMSO in 100 μl of culture medium for 4 days. Then, PDOs were collected and subjected to proteomics assay, quantitative analysis of energy metabolism, immunofluorescence staining and RT-qPCR

For immunofluorescence staining of PDOs, the medium was carefully aspirated after 4 days incubation with gamitrinib or DMSO, and 100 μl of live/dead reagents (L3224, Invitrogen) was added followed by 20-30 min of incubation at room temperature in the dark. Images were acquired with imaging system (IX73, OLYMPUS, Japan).

### RT-qPCR of PDOs

Total RNA was extracted from PDOs. 1 μg of total RNA was reverse transcribed into complementary DNA (cDNA) using PrimeScript™ RT Reagent Kit (Takara, RR047B). Amplification of cDNA product was performed using specific primers with the TB Green® Premix Ex Taq™ II (Takara, RR820B) on a Real-Time PCR detection system (BioRad). Samples were analyzed in triplicate, and β-tubulin levels were used for normalization. Primers sequences for RT-qPCR were as follows: SDHB: forward primer, 5’-AAGCATCCAATACCATGGGG-3’, and reverse primer, 5’-TCTATCGATGGGACCCAGAC-3’; COX7B: forward primer, 5’-CTTGGTCAAAAGCGCACTAAATC-3’, and reverse primer, 5’-CTATTCCGACTTGTGTTGCTACA-3’; ATP5B: forward primer, 5’-CAAGTCATCAGCAGGCACAT-3’, and reverse primer, 5’-TGGCCACTGACATGGGTACT-3’; and β-actin: forward primer, 5’-GAGAAAATCTGGCACCACACC-3’, and reverse primer, 5’-GGATAGCACAGCCTGGATAGCAA-3’.

### Animal experiments

All studies were approved by the Committee for Animal Use and Welfare at Guangzhou Medical University (protocol 2021169). C57BL/6 female mice at the age of 6-8weeks were purchased from GemPharmatech Co.Ltd. Animals were housed under standard vivarium conditions (22±1 °C; 12 h light/ dark cycle; with ad libitum food and water). For anesthesia, mice were first anesthetized in 5% Isoflurane, and then maintained on 1.5–2% throughout the procedures. When anesthetized, core body temperature of animals was maintained at 37°C. LLC cells expressing luciferase tag (LLC-Luc) were cultured in complete media (RPMI1640 supplemented with 10% fetal bovine serum and 1% penicillin/streptomycin). On the day of experiment, cells were washed and harvested with 0.25% trypsin-EDTA (ThermoFisher) when ∼90% confluence was reached. Cells were centrifuged at 80 × g for 3 min. Subsequently, cells were washed in serum-free media twice to remove residual serum, counted with a hemocytometer and re-suspend in Hank’s Balanced Salt Solution (HBSS) at the density of 1-5 × 10^6^ cells/ml. Throughout the injection period, cell suspension was kept on ice until time of injection. The experiment was completed within 3 hours of cell harvesting and more than 95% of cells were viable. After a habituation period of one week, mice were administered 5% isoflurane for anesthesia induction and 1.5% isoflurane for anesthesia maintenance in 30% O_2_/70% N_2_O through a face mask. Each mouse was placed in supine position and fixed on an operating table. The middle of the neck was sterilized and one 1cm incision was made to expose the trachea. The left side of the trachea was separated from the muscle to expose the carotid sheath. Extra attention was paid to protect the blood vessels and minimize bleeding. The common carotid artery was bluntly separated from the vagus nerve under a stereo microscope, and an 8-0 silk suture was placed on the common carotid artery to separate the common carotid artery. The bifurcation, and the external carotid artery distal to the bifurcation was ligated with silk thread to block the blood flow of the external carotid artery. The proximal segment of the common carotid artery was temporarily blocked with a small vessel clip. Tumor cell suspension was injected as follows: resuspend cells stored on ice, draw 100 μl with a 100 μl Hamilton microsyringe, carefully remove any air bubbles, and puncture the common carotid artery under a microscope. The injection was completed within 1 min. After the cell suspension entered the blood vessels, the color of the nearby blood vessels and muscle tissue was pale under the microscope, confirming that the cell suspension successfully entered the carotid system. After the injection was completed and the needle of the syringe was withdrawn, the slip knot at the distal end was lifted quickly. At that time, the ligature was deliberately loosened slightly to ensure that any bubbles that may enter the blood vessel cavity may be discharged at the injection site. We ligated the distal end of the common carotid artery by tying the knot with both hands, and the skin was sutured. The entire procedure was completed within approximately 15 minutes by fully trained personnel. One week after injection of tumor cells, in vivo bioluminescence was performed to confirm the tumor formation of engrafted LLC-Luc cells. Mice were anesthetized and retro-orbitally injected with luciferin (150 mg/kg; PerkinElmer cat. # 122799,). Images were acquired by using NightOWL II LB 983 In Vivo Imaging System (Berthold Technologies GmbH, Bad Wildbad, Germany) in order to measure the bioluminescent activity of the luciferase enzyme. Fixed-area region of interests (ROIs) were created overhead, and photons emitted from the ROIs were quantified. After tumor formation detected by bioluminescence, mice were randomized to receive treatments and the investigators were not blinded. Mice were injected intraperitoneally with vehicle, gamitrinib (MedChemExpress cat.# HY-102007A) at 10 mg/kg every day, anti-PD-1 antibody (BioXcell cat.# BE0146) at 100ug per mouse every 3 days, or the combination of gamitrinib plus anti-PD-1 antibody administered at the same dose. The endpoint of experiment was a moribund state of the animal in accordance with the Institutional Animal Welfare Regulations. At the end of the experiment, mice were euthanatized and brains were harvested and analyzed by histology. OS of mice was defined as the time between the beginning of treatment and euthanasia of mice.

## Data availability

All raw RNAseq and WES data have been deposited in the National Genomics Data Center. Access to the data can be requested by completing the application form via https://ngdc.cncb.ac.cn/gsa-human/ (accession number: HRA004247 for WES data and HRA005036 for RNAseq data).

## Acknowledgments

The authors thank all collaborators for helpful comments and discussions, and thank Accurate International Biotechnology Co. for their assistance with the organoid techniques. The authors also thank Li Li, Chunjuan Bao and Fei Chen from the Institute of Clinical Pathology of West China Hospital for the technical support in histological staining. This work was supported by the National Natural Science Foundation of China (81872324 to YG. Mou, 31771549 to Y.C., and 82203574 to S.Wei), the GuangDong Basic and Applied Basic Research Foundation (2020A1515110069 to H.D.), and the Science and Technology Support Program of Sichuan Province (23ZDYF2661 to S.Wei).

## Author contributions

H.D., L.C., Y.C., G.Z., L.Liu. and YG.Mou. conceived the original idea, designed the study, and provided financial and administrative supports. Z.Wei., Y.C., G.Z., L.Liu., YG.Mou, M.Y., and L.C. jointly supervised the study. S.Wei., C.L., YG.Mou., L.C., X.Z. and C.G. provided study materials or patients, and contributed to clinical data interpretation. H.D., S.Wei., C.L., M.L., S.Wu, W.H., L.Liang, C.Y., and YH.Mou. participated in collection of clinical samples and data. J.R., Z.Y. and Z.Wei. contributed to data analysis and data interpretation. S.Wei, C.L. and Y.J. performed staining of FFPE slides. H.D., Z.Wang and M.Y. performed animal experiments. Y.L., H.L., Y.X., Z.C., E.S. and R.D. contributed to data interpretation and manuscript edit. All authors involved in writing the manuscript and final approval of the manuscript.

## Supplemental Figure Legends

**Supplemental Figure 1. Related to Figure 1**. Analysis pipeline for whole-exome sequencing (WES), RNA sequencing (RNAseq), single cell RNA sequencing (scRNAseq), proteomics, and metabolomics data. Flow-chart demonstrates sequential use of tools in evaluation of DNA, RNA, single cell RNA, protein and metabolite samples.

**Supplemental Figure 2. Related to Figure 2. a**. Tumor mutation load comparison in primary, brain metastasis (BrM) against 33 TCGA cohorts derived from MC3 project. Top annotation is number of samples in each cohort, and y-axis is the logscale of tumor mutational burden with median mutation marked with red horizontal line. **b**. Box plots to show mutational burden comparison in matched primary lung cancer and BrM. The p value was calculated by using Wilcoxon tests.

**Supplemental Figure 3. Related to Figure 3. a.** Oncoplot showing the specifically mutated genes in paired primary lung cancer and lung cancer brain metastasis (LC-BrM) specimens. **b.** Forrest plot to show the specifically mutated genes comparison in paired primary lung cancer and LC-BrM tumor specimens. **c and d.** Cluster plots of the primary lung cancer (left) and LC-BrM sample (right) derived from patient DHP7 (**e**) and Lung-10 (**f**). X-axis represented variant allele frequency, the top bar showed the number of clusters on top of each plot, and the math number was noted in the upper left corner.

**Supplemental Figure 4. Related to Figure 4. a and b.** Copy number gain (**a**) or loss (**b**) on the primary lung cancer (left column) and matched brain metastasis (BrM) samples (right column). **c and d**. Heatmap of copy number profiles for samples from primary lung cancer (**c**) and BrMs (**d**). Each row represents the copy number profile of a tumor sample across chromosomes 1 to 22 and X. Red indicates copy number gain; blue, loss. e and f. Specific genes of copy number gain and loss in primary lung cancer (**e**) and BrMs (**f**). Red indicates copy number gain (CN >= 3, CN2-3); yellow, no change (copy neutral); blue, loss (Deletion, Loss CN1).

**Supplemental Figure 5. Related to Figure 5. a and b.** PCA plots to show the distribution of samples from two cohorts of 56 patients before (**a**) and after (**b**) ComBat process. **c.** The table of top 20 ranked gene pathways that were positively correlated with the lung cancer brain metastasis (LC-BrM) phenotype. Pathways were ordered by p-value and normalized enrichment score (NES). Five pathways related to mitochondrial biogenesis and oxidative phosphorylation were highlighted with different colors. **d.** tSNE plot of ∼20,000 single epithelial cells with color-coded by two major cell lineages: 15,463 mbrain (brain metastasis) single cells and 7,270 tLung (primary lung cancer) single cells. **e-l**. tSNE plots and box plots of enrichment score of single cells for the rest four mitochondrial pathways mentioned in Figure 5b. **m**. The box plot of enrichment score of the citric acid TCA cycle pathway in primary lung cancer and brain metastasis from spatial transcriptomic data (Zhang et al. 2022). **n**. Relative mitochondrial DNA (mtDNA) content of paired primary lung cancer and brain metastasis lesions. The p-value was determined using pairwise two-sided Student’s t test. **o.** The table of top 20 ranked protein pathways that were positively correlated with the LC-BrM phenotype. Pathways were ordered by p-value and NES. Four pathways related to mitochondrial biogenesis and oxidative phosphorylation were highlighted with different colors. **p.** Box plots showing the protein expression of MTCO1, UQCRC2 and COXIV based on reverse phase protein array data in nine paires of primary lung and brain metastasis lesions. The p-value was determined using pairwise two-sided Student’s t test. **q.** Box plots for the immunohistochemistry scores of MTCO1, UQCRC2 and COXIV in 51 paires of primary lung and brain metastatic lesions. The p-value is determined using pairwise two-sided Student’s t test. *, p<0.05; **, p<0.01; ***, p<0.001; ****, p<0.0001.

**Supplemental Figure 6. Related to Figure 6. a.** The table of bottom 20 ranked gene pathways that were negatively correlated with the phenotype of lung cancer brain metastasis (LC-BrM). Pathways were ordered by p-value and normalized enrichment score (NES). Five immune related pathways were highlighted with different colors. **b and c.** Box plot representation of xCell (**b**) and CIBERSORTx (**c**) scores for patients with paired primary lung cancer and LC-BrMs. The p*-*value was determined by use of pairwise t-test. **d.** tSNE plot of ∼20,000 lymphocytes with color-coded by two major cell lineages: 3,910 mbrain (brain metastasis) single cells and 18,587 tLung (primary lung cancers) single cells. **e-n.** tSNE plots and box plots of enrichment score of single cells for the five immune pathways mentioned in Figure 6a. **o**. The table of bottom 20 ranked protein pathways that were negatively correlated with the phenotype of LC-BrM. Pathways were ordered by p-value and NES. Four immune related pathways were highlighted with different colors.

**Supplemental Figure 7. Related to Figure 7. a.** The heatmap showing the enrichment of indicated five mitochondrial pathways and five immune pathways in an independent cohort of 63 brain metastasis (BrM) lesions with mRNA microarray data (Pocha et al. 2020). **b.** Scatter plot showing the IC_50_ to gamitrinib of 20 BrM patient-derived organoids.

**Supplemental Figure 8. Related to Figure 8. a-e.** Bar plots of quantification analyses of immunohistochemistry staining of Ki-67 (**a**), CD3 (**b**), CD4 (**c**), CD8 (**d**), and PD-L1 (**e**) in Lewis lung carcinoma brain metastases. The p-value is determined using two-sided Student’s t test. ***, p<0.001; ****, p<0.0001.

