## Supplemental Figures for "Integrated Analyses of Multi-omic Data Derived from Paired Primary Lung Cancer and Brain Metastasis Reveals the Metabolic Vulnerability as a Novel Therapeutic Target"

### Supplemental Figure 1

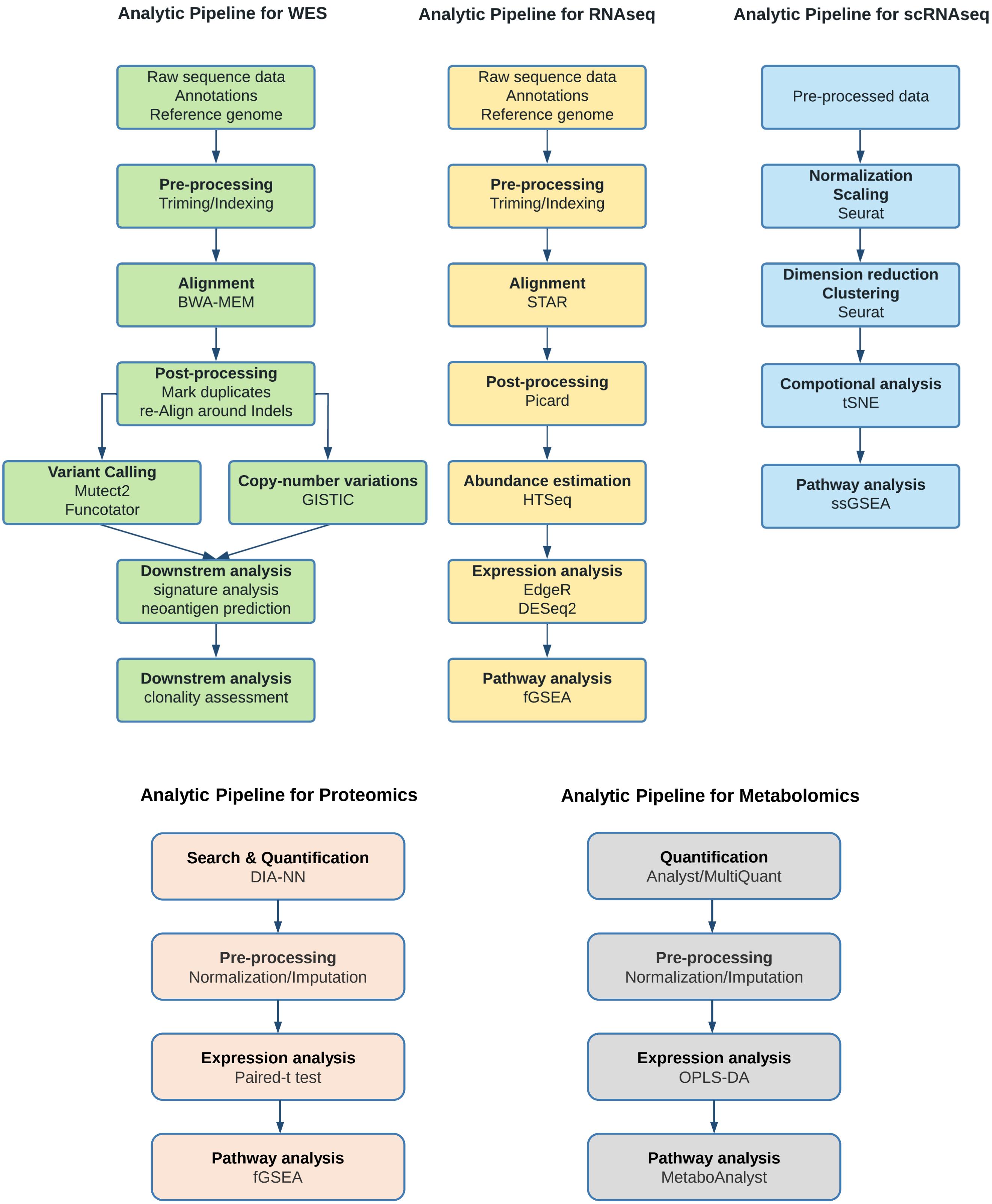

### Supplemental Figure 2

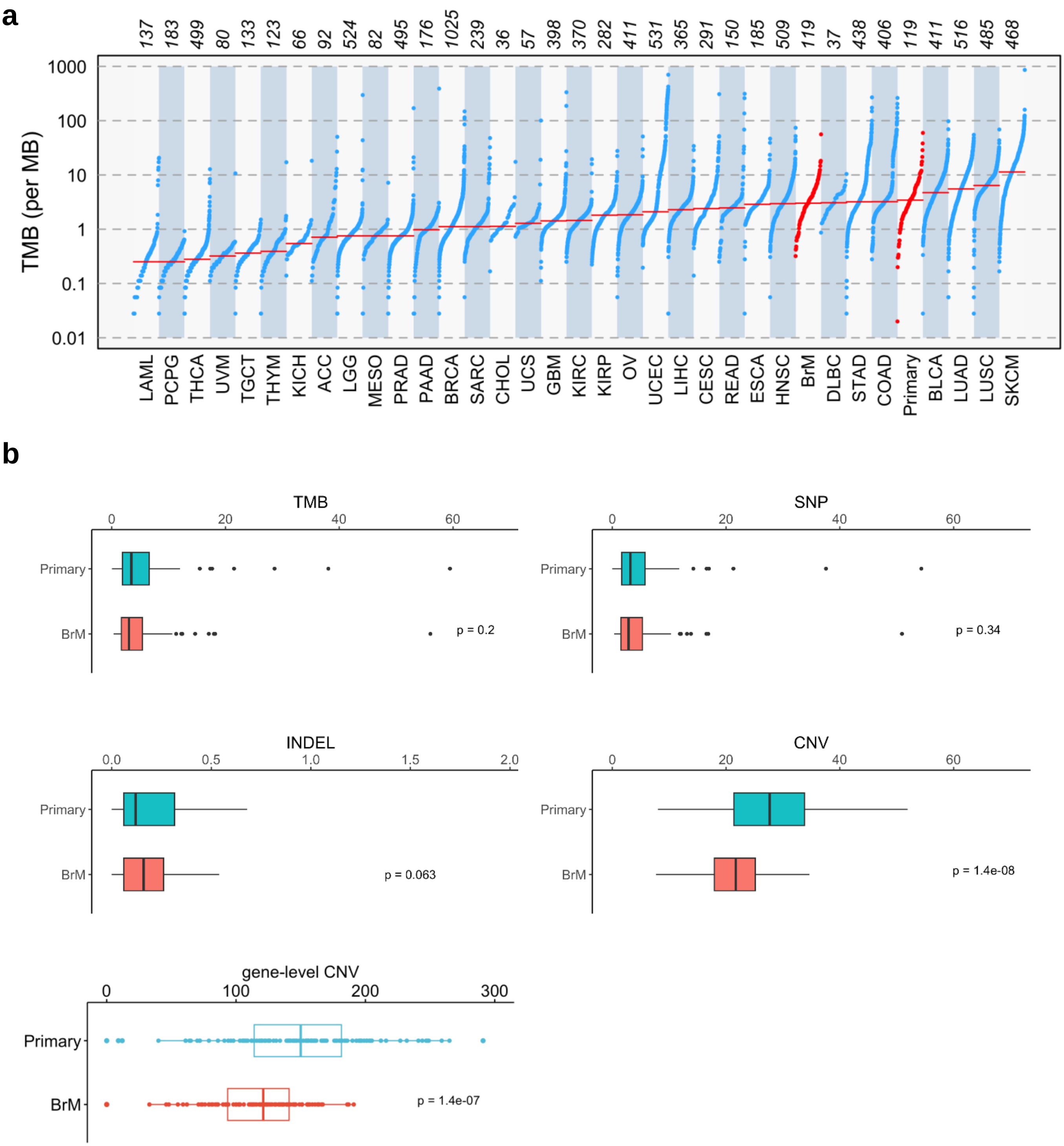

### Supplemental Figure 3

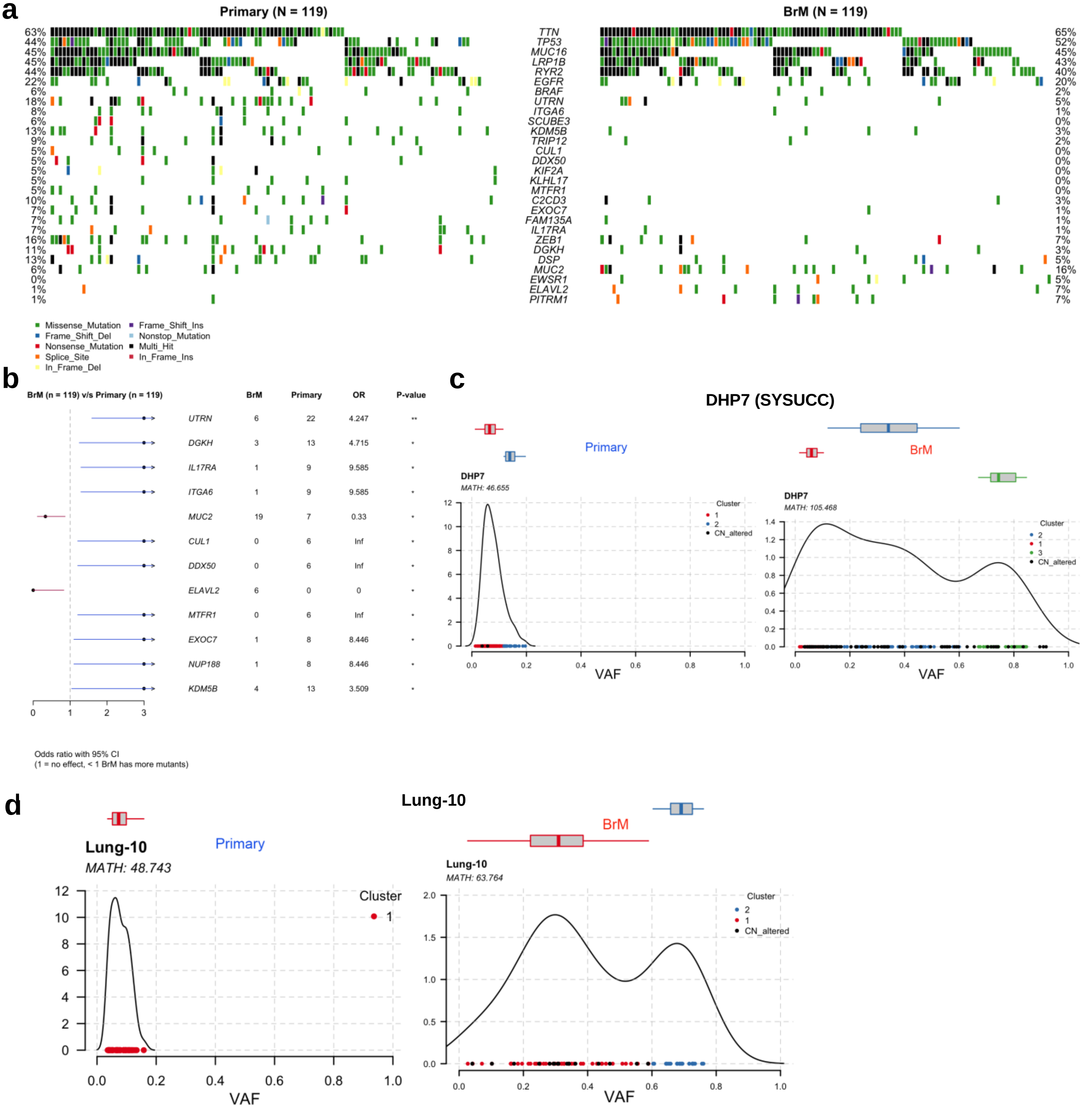

### Supplemental Figure 4

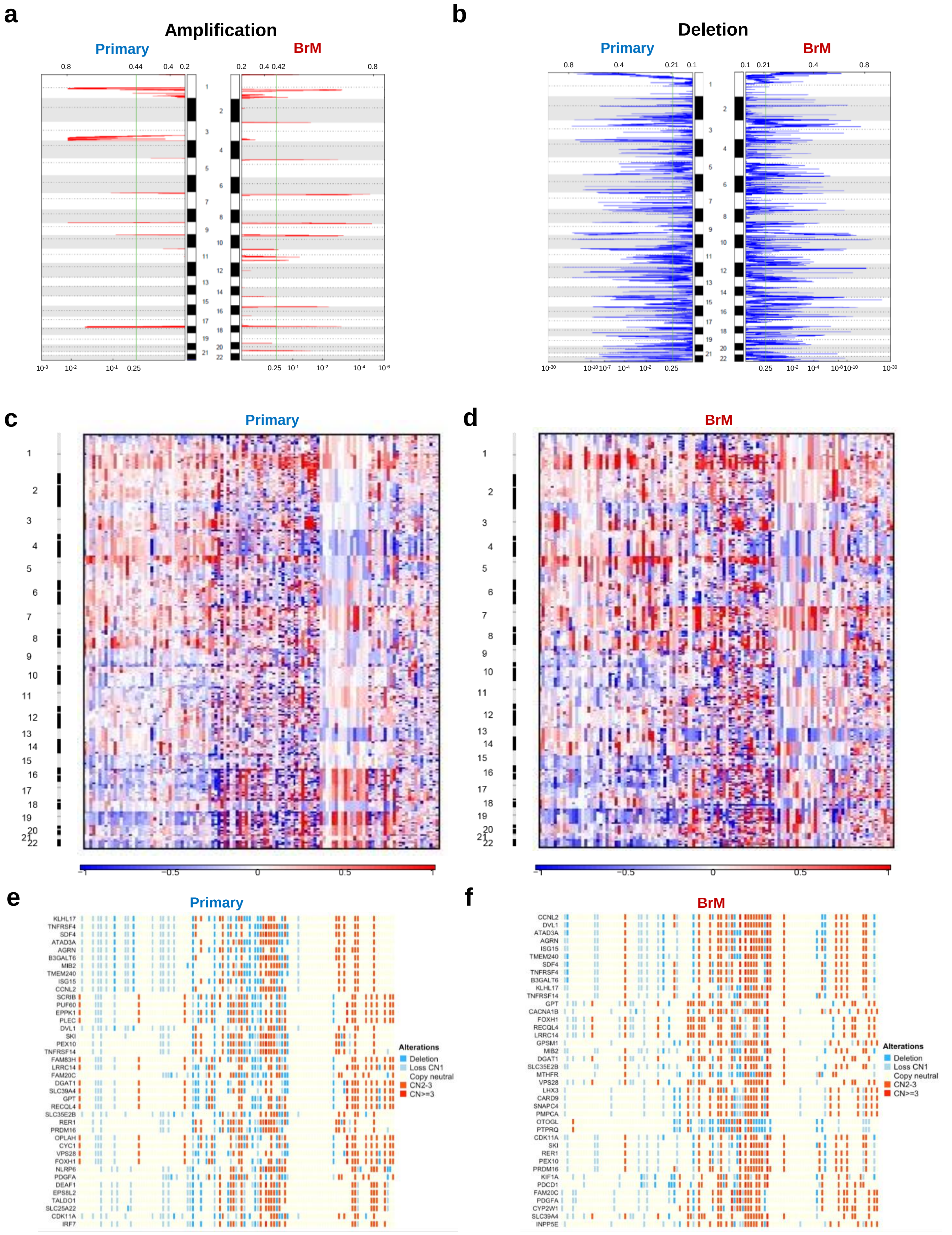

### Supplemental Figure 5

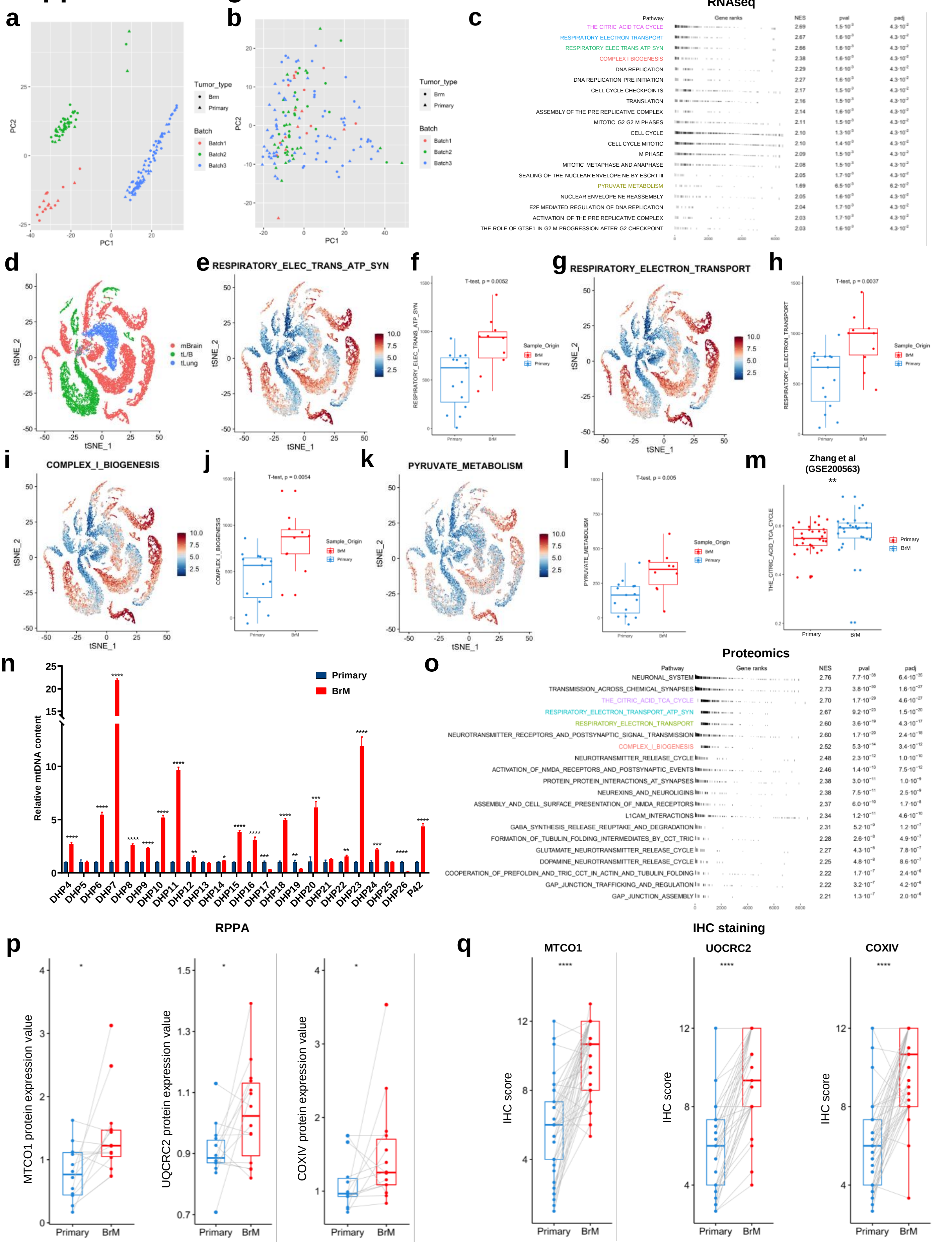

### Supplemental Figure 6

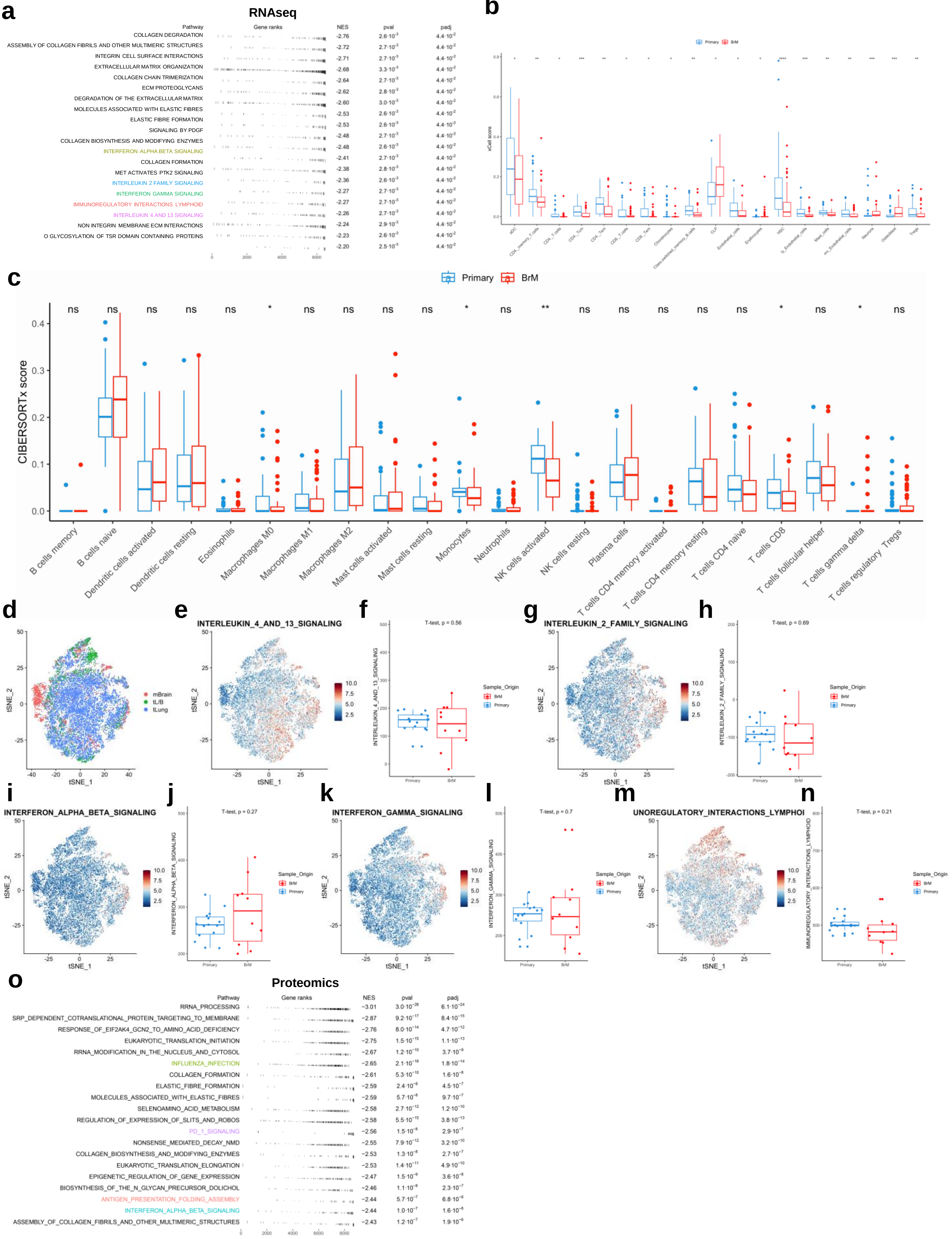

### Supplemental Figure 7

**a**

Pocha et al. 2020

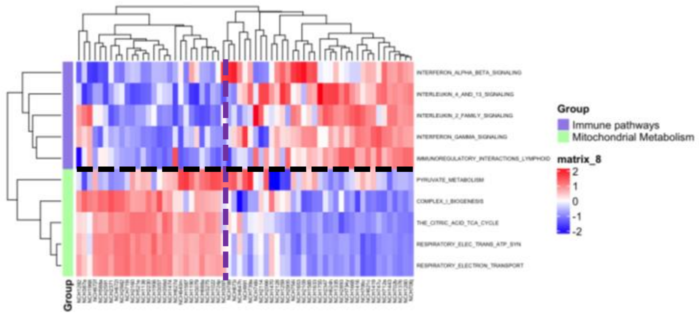


**b**

PDOs of LC-BrM (N=20)

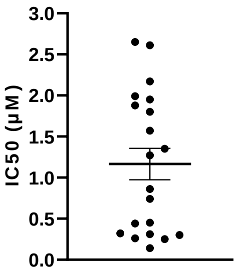

### Supplemental Figure 8

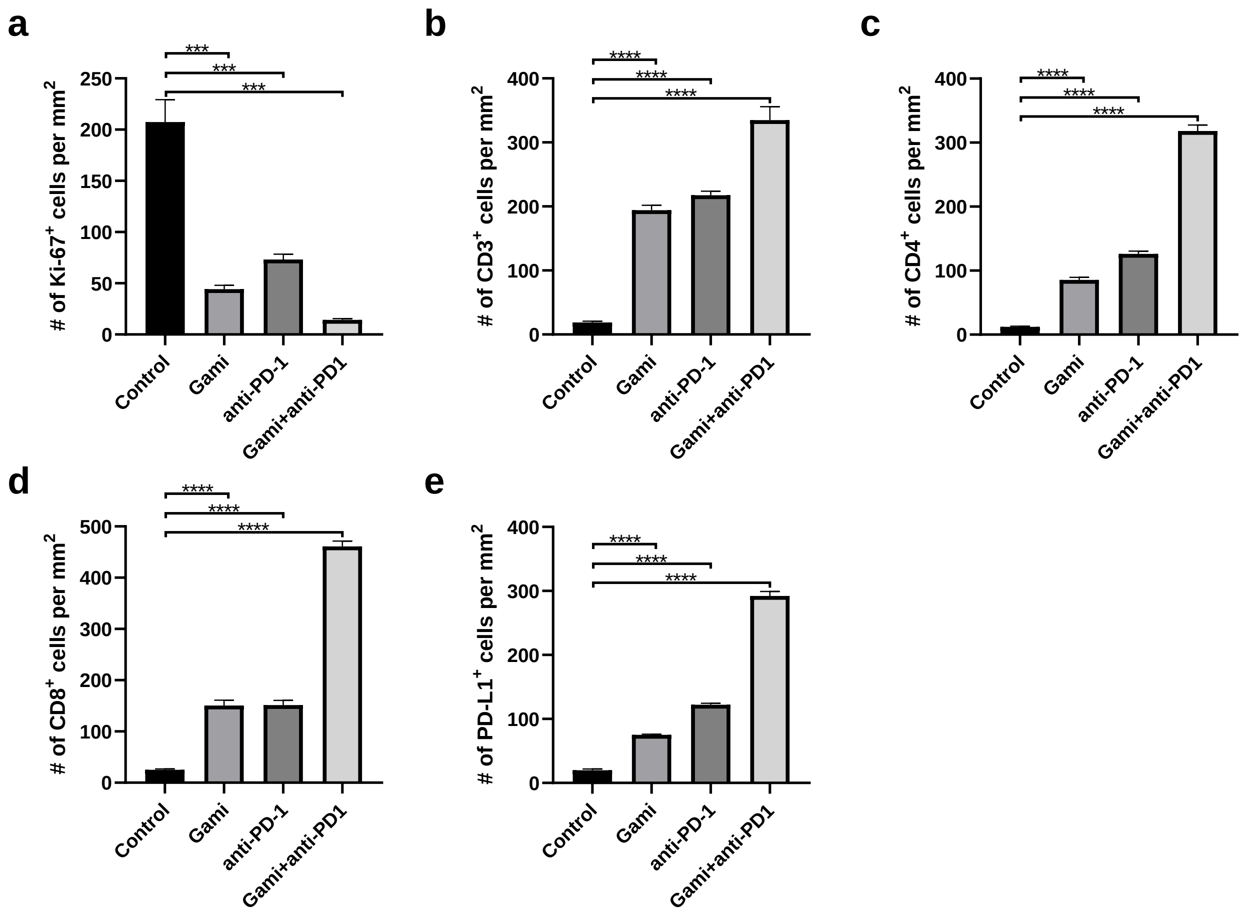
